# Computational mechanisms of curiosity and goal-directed exploration

**DOI:** 10.1101/411272

**Authors:** Philipp Schwartenbeck, Johannes Passecker, Tobias U Hauser, Thomas H B FitzGerald, Martin Kronbichler, Karl Friston

## Abstract

Successful behaviour depends on the right balance between maximising reward and soliciting information about the world. Here, we show how different types of information-gain emerge when casting behaviour as surprise minimisation. We present two distinct mechanisms for goal-directed exploration that express separable profiles of active sampling to reduce uncertainty. ‘Hidden state’ exploration motivates agents to sample unambiguous observations to accurately infer the (hidden) state of the world. Conversely, ‘model parameter’ exploration, compels agents to sample outcomes associated with high uncertainty, if they are informative for their representation of the task structure. We illustrate the emergence of these types of information-gain, termed active inference and active learning, and show how these forms of exploration induce distinct patterns of ‘Bayes-optimal’ behaviour. Our findings provide a computational framework to understand how distinct levels of uncertainty induce different modes of information-gain in decision-making.

## Introduction

The balance between *exploitation*, i.e. choosing the most valuable option given current beliefs about the world, and *exploration*, i.e. choosing options that allow us to forage and learn about our environment, lies at the heart of decision-making and adaptive behaviour (Cohen, McClure, & Yu, 2007). The trade-off between choosing to exploit or explore is a key focus of computational theories of behaviour in both artificial intelligence and neuroscience, such as in reinforcement learning and Bayesian models of behaviour (Friston et al., 2015; Friston et al., 2017; Sun, Gomez, & Schmidhuber, 2011; Sutton & Barto, 1998; Yang, Wolpert, & Lengyel, 2016; Houthooft et al., 2016). Importantly, recent behavioural evidence suggests that humans perform a mixture of both *random* and *goal-directed* exploration (Gershman, 2018a, 2018b; Wilson, Geana, White, Ludvig, & Cohen, 2014). Random exploration has been introduced in early accounts of exploratory behaviour (Daw, O’Doherty, Dayan, Seymour, & Dolan, 2006; Sutton & Barto, 1998). This behaviour is defined as a deviation from the currently most valuable policy by randomly sampling any other option. A classical way of formalising random exploration is via ε-greedy or softmax choice rules, where in the latter the tendency towards randomness is governed by an inverse temperature parameter (Sutton & Barto, 1998). A more refined account of random exploration has been introduced via Thompson sampling (Thompson, 1933), where an agent samples from a posterior over reward statistics and chooses the most valuable option with respect to this sample, thus taking its uncertainty over reward statistics into account (Agrawal & Goyal, 2011; Speekenbrink & Konstantinidis, 2015).

In contrast to random exploration, goal-directed, information-seeking exploration is guided by the uncertainty in an agent’s model of the world and motivates curiosity-driven behaviour (Gottlieb, Oudeyer, Lopes, & Baranes, 2013). This implies that agents will selectively sample options that are informative, i.e. that are associated with the highest uncertainty. A prominent example of uncertainty-sensitive exploration is the upper confidence bound algorithm (Agrawal, 1995; Auer, 2002; Kaelbling, 1994; Sutton & Barto, 1998), which adds an uncertainty bonus (Kakade & Dayan, 2002) to options that have not been sampled for a long time or that are associated with high uncertainty. See (Gershman, 2018a, 2018b) for a discussion of these two types of exploration and specific predictions arising from these formulations, which we will discuss in more detail below.

It is challenging to provide a formal account of the trade-off between behaviour that aims at maximising reward and fulfils an agent’s preferences over states on the one hand and acquiring information about the world on the other. Furthermore, an important challenge lies in moving beyond descriptive accounts of behaviour towards understanding the generative mechanisms of information gain that could be implemented by a biological system. A particularly challenging aspect lies in providing a formal account of goal-directed exploration, where agents are guided by minimising uncertainty and actively learning about the world. This is particularly delicate because one can dissociate different types of uncertainties. For example, if I offered you an option that may have a positive or a negative outcome, I leave you in a state of uncertainty at two levels. First, you have no idea about the probabilities of winning or losing. For example, there could be a 50% or 99% chance of winning. Second, even if you knew the probability of winning exactly, there will still be some uncertainty about the outcome if you chose the option. These types of uncertainties have been termed unexpected and expected uncertainty (Yu & Dayan, 2005) or, in economics, ambiguity and risk. The key point is that it is necessary to resolve ambiguity first before agents can assess the value of options and their associated risk.

We discuss these different aspects of behaviour in terms of active Bayesian inference, by casting choice behaviour and planning as variational probabilistic inference (Friston et al., 2013; Friston, Fitzgerald, Rigoli, Schwartenbeck, & Pezzulo, 2017). Here, agents are assumed to form expectations over observable states (outcomes) and infer policies that minimise the expected information-theoretic surprise about these observations. These expectations reflect an agent’s preferences over observations. Thus, by minimising surprise, agents find policies that make visiting preferred states more likely. This information-theoretic quantity can be approximated by the expected free energy, which is a function of (approximate posterior) beliefs about the states of the world, formed under a generative model based on a Markov decision process, as will be described below.

Under this approach, different types of exploitative (*pragmatic*) and exploratory (*epistemic*) behaviour emerge. The key aspect that motivates goal-directed uncertainty reduction is the mapping from (hidden) states to observations. This form of uncertainty reduction becomes relevant in partially observable problems, where in addition to inferring the best policy; agents also have to infer the current (hidden) state that caused an observation. In order to minimise uncertainty about the current state, agents can try to navigate to (observable) outcomes, where the mapping to the underlying hidden state is unambiguous. A simple example is a bird that is searching for prey: in the case of high uncertainty about the prey’s location, a bird might go to a vantage point first to minimise uncertainty about the prey’s location (i.e., the underlying hidden state), before predation. Another example is contextual inference, where agent’s need to disclose the current context (i.e., the hidden state), in order to infer what to do (e.g., is there milk in the fridge?). In case of contextual uncertainty, agents will prefer to sample outcomes that allow for precise inference about the current context, before making a choice about where to look for reward. Formally, this means that agents will try to actively sample outcomes that have an unambiguous (low conditional entropy) mapping to hidden states – hence *active inference* allowing for ‘hidden state exploration’.

Importantly, the exact same ‘epistemic’ imperatives apply to beliefs about model parameters that describe a subject’s knowledge about state transitions or the probability of various outcomes given the underlying (hidden) states. In other words, uncertainty about states of the world is accompanied by uncertainties about the lawful contingencies that underwrite state transitions and the relationship between hidden states and observable outcomes. In contrast to the examples above, which reflect uncertainty about the underlying hidden state, given an agent’s model of the task, this form of uncertainty reflects an agent’s ignorance about the causal structure of the model per se. For example, agents can be uncertain about the current context that determines the value of options (i.e., uncertainty about a hidden state) or uncertain about the value of options given a current context (i.e., uncertainty about model parameters). To reduce the latter type of uncertainty, agents can expose themselves to observations that complete ‘knowledge gaps’ and thereby enable learning about the probabilistic structure of unknown and unexplored (novel) contingencies – hence *active learning* allowing for ‘model parameter exploration’.

In the following, we will introduce the theoretical framework underlying active inference and active learning and use simulations to illustrate the emergence of these particular types of exploratory behaviour. We will consider the resolution of uncertainty about *states* and *parameters* in terms of *salience* and *novelty* respectively; where ‘salience is to inference’ as ‘novelty is to learning’. We will use a simple two-armed bandit problem in which a subject has to choose between a risky high reward and a safe low reward, where the probabilities of the risky option are unknown. Minimising expected free energy leads to curiosity-driven *active learning* that initially favours the novel risky option – because this option provides uncertainty reduction about an agent’s parameterisation of the task. We will also show how the same computational framework motivates *active inference* in situations where certain actions disclose salient information about hidden states, such as whether there is currently a high or low reward probability in the risky option. Based on this paradigm, we will illustrate different sorts of explorative behaviour, contrast them with random exploration or purely exploitative choices, and consider how different tendencies emerge under different priors over beliefs about outcomes and the precision of those beliefs.

## Theoretical Framework: Probabilistic inference and free energy

Our theoretical approach assumes that agents, such as brains or economists, minimise the expected free energy of future outcomes and hidden states (Friston, 2013). This premise allows to derive generic update rules for action (i.e., policy selection), perception, and learning based on variational Bayes, which is described briefly in this section.

Active inference rests upon a generative model of observed outcomes. This model is used to infer the most likely causes of outcomes in terms of expectations about states of the world. These states are called *hidden* because they are usually not or only partially observable and can only be inferred through observations. Importantly, agents can also infer different actions that determine the most likely observations they will make. This means that observations depend upon action, which requires the generative model to infer expectations about counterfactual outcomes under different actions or policies. Given that the generative model enables inference about hidden states based on observations, agents can also form expectations about future states. The ‘optimisation’ of these expectations (i.e., state estimation) is cast as minimising variational free energy, which finds the most likely (posterior) expectations about states of the world, given current observations. In addition to forming posterior beliefs about hidden states, active inference requires posterior expectations about the policy or action sequence currently being pursued. Crucially, the prior probability of a policy decreases with the free energy expected under that policy. This (expected) free energy is a proxy for surprise or model evidence, and thus allows to cast choice behaviour as minimising expected surprise or uncertainty (or, equivalently, maximising expected model evidence) (Friston et al., 2015). This provides a formal grounding for the notion of the ‘value’ of a policy; such that the value is defined with respect to an agent’s generative model of the world, and valuable policies maximise the expected log-evidence of that model, a process sometimes referred to as ‘self-evidencing’ (Hohwy, 2016).

### The generative model

Agents are assumed to perform approximate inference based on variational Bayes, which casts a difficult and usually intractable inference problem as a bound optimisation problem (Beal, 2003; Bogacz, 2017). This implies that expectations about hidden states are updated to minimise variational free energy under a generative model. Figure 1 provides the specification of the Markovian generative model used in the simulations below. Outcomes (observations) at a particular discrete time-step depend upon true hidden states in the world, while hidden states evolve according to Markovian transition probabilities contingent upon actions emitted by an agent. The generative model is specified by two sets of arrays. The first, ***A***, maps from hidden states to outcomes. That means that ***A*** models an agent’s observation model or the emission function in a hidden Markov model, specifying the likelihood of an observation under a given hidden state. The second, ***B***(*u*), prescribe the transitions among hidden states, given an action, *u*. These transitions are Markovian, such that the probability of the subsequent state is fully determined by the current state and action. The remaining parameters encode prior expectations (i.e., preferences or utilities) about observations, ***c***, and initial states, ***d***.

**Figure 1.**
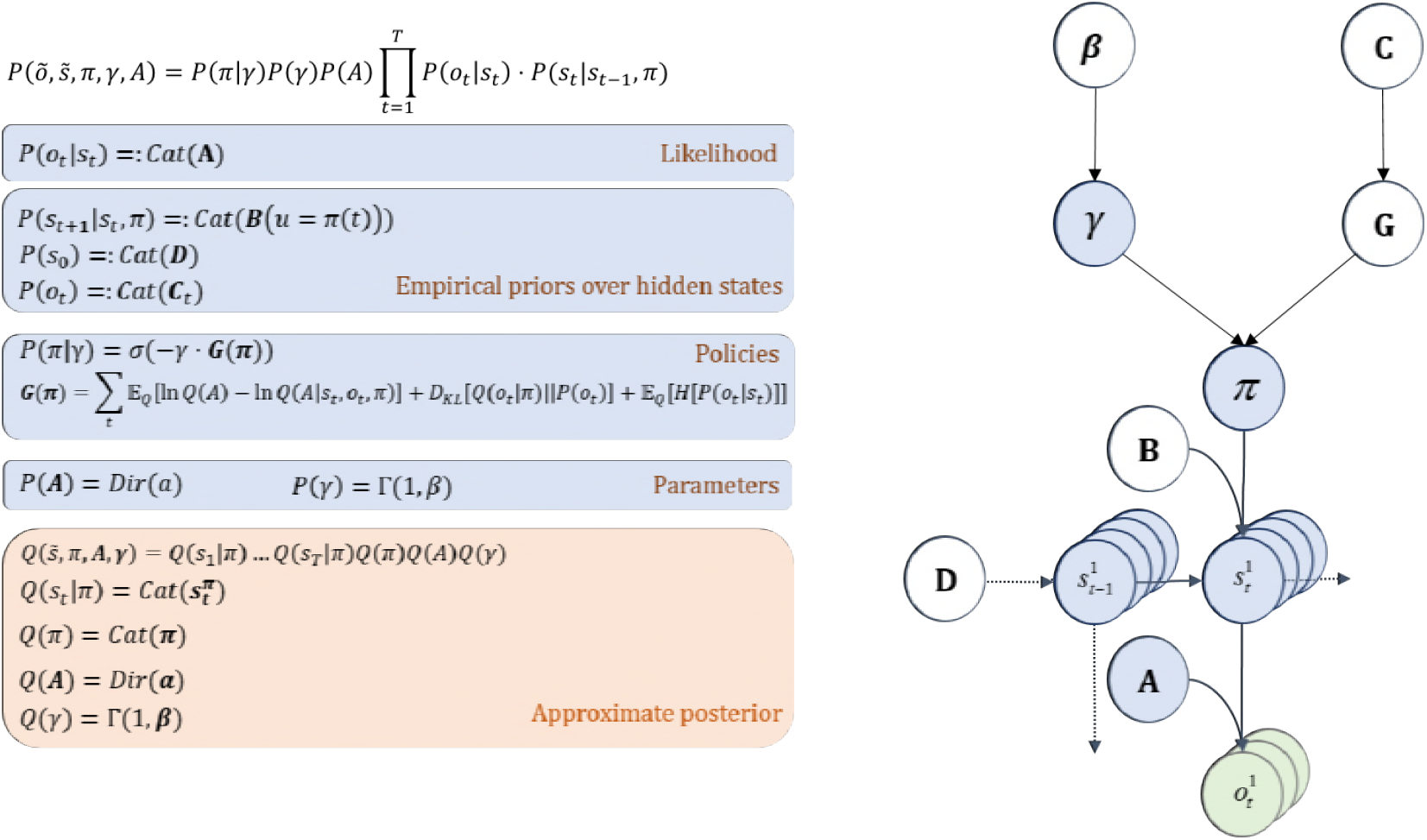
Generative model and approximate posterior. A generative model specifies the joint probability of observations and their hidden causes. The model is expressed in terms of a *likelihood* (the probability of observations given causes) and priors over causes. Here, the likelihood is specified by a matrix *A* whose components are the probability of an outcome under all possible hidden states, *P*(*o*_*t*_|*s*_*t*_). The empirical priors in this instance pertain to transitions among hidden states *B* that depend upon action, *P*(*s*_*t*+1_|*s*_*t*_. *π*), where actions are determined probabilistically in terms of policies (sequences of actions, *π*). The key aspect of this generative model is that policies are more probable *a priori* if they minimise the (sum or path integral of) expected free energy ***G***(*π*). Approximate inference on the hidden causes (i.e., the current state, policy, precision and model) proceeds using variational Bayes. In variational Bayesian inference (*model inversion*), one has to specify the form of an approximate posterior distribution, which is provided in the lower panel. This form uses a mean field approximation, in which posterior beliefs are approximated by the product of marginal distributions over hidden causes. The figure on the right shows the directed graphical model of the dependencies implied by the equations on the right. Cat = categorical distribution, dir = Dirichlet distribution, Γ = Gamma distribution.

The posterior mapping from hidden states to outcomes **(*A*)** or hidden states **(*B*)** are parameterised as Dirichlet distributions, whose sufficient statistics are concentration parameters (Friston et al., 2016). These concentration parameters effectively reflect the (normalised) number of times a particular combination of states and outcomes has been encountered. In the following simulations on active learning, we focus on learning the observation model and will assume that state transitions and observations/initial states are known or fixed. How the state space and the dimensions of the different matrices that determine the mapping between these states are themselves learned is an important and interesting question but goes beyond the scope of this paper (see discussion).

The generative model illustrated in Figure 1 implies that outcomes (observations) are generated in the following way: first, a policy is selected using a softmax function of expected free energy for each policy (see below), which also depends on an agent’s degree of randomness (precision) in behaviour. Sequences of hidden states are then generated based on the probability transitions specified by the selected policy. These hidden states then generate outcomes. Perception (state inference) corresponds to inverting the generative model given a sequence of outcomes, while (parameter) learning corresponds to updating the mapping between hidden states and outcomes. Consequently, ‘perception’ corresponds to inferring or optimising expectations about hidden causes with respect to variational free energy, while learning corresponds to accumulating concentration parameters. These variables constitute the sufficient statistics of the approximate posterior beliefs, denoted by the probability distribution *Q*(*s, π, A, γ*), where *s, π, A, γ* are the hidden or unknown variables.

### Variational free energy and inference

Having specified a Markovian generative model and the approximate posterior, the last step is to define the variational free energy and resulting update equations that are used to infer hidden causes, and the expected free energy over future states under policies, which defines the value of a policy.

Variational Bayesian inference implies that by minimising variational free energy with respect to the specified posterior *Q*(*x*) over hidden causes *x* (where *x* = {*s, π, A, γ*} in our example) we approximate the true posterior 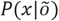:

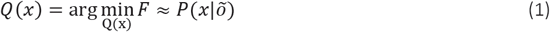

There are several equivalent expressions for variational free energy: one is in terms of the entropy minus energy:

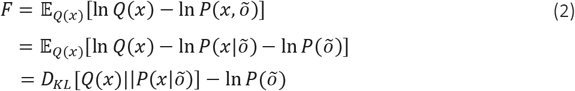

where 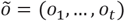 denotes observations up until the current time *t*. Because the (KL) divergence cannot be less than zero, the last equality means that free energy is minimised when the approximate posterior *Q*(*x*) becomes the true posterior 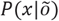. In this case, the variational free energy becomes the negative log evidence for the generative model (Beal, 2003).

Rewriting equation (1) shows that variational free energy can also be written as

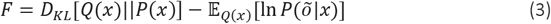

This implies that minimising variational free energy maximises the expected likelihood of observations under the approximate posterior (‘accuracy’) whilst minimising the divergence between the approximate and true distribution over hidden causes (‘complexity’). Having defined the objective function, the sufficient statistics encoding posterior beliefs can be updated by minimising variational free energy, as discussed in detail in (Friston et al., 2017; see also appendix of Parr & Friston, 2018 for the derivation of these updates). Given the focus of this paper, we will discuss inference on valuable policies in detail below.

As we have shown above, minimising free energy ensures that expectations about hidden causes are close to the true posterior over hidden causes, given observed outcomes. However, if we want to apply this notion to define the value of actions and policies, we need to consider potential *future* outcomes and states under a given policy. This can be achieved by making the log prior probability of a policy the (negative) free energy expected under that policy (Friston et al., 2017):

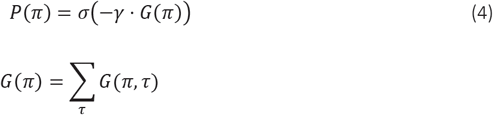

where *τ* refers to a time-step in the future, *τ* ∈ {*t* + 1, …, *T*} with *t* reflecting the current time step. Note that the expected free energy over future states that determines the value of a policy resembles the expected value of future reward in reinforcement learning (Sutton & Barto, 1998), although there is no discount parameter over future states. *γ* reflects a precision parameter that governs an agent’s goal-directedness and randomness in behaviour, parameterised by a gamma function with rate parameter *β* (see Figure 1). Based on these beliefs about policies, agents sample an action, where the randomness of this sampling is governed by a precision parameter *α*. The parameters *β* and *α* can be thought of as inverse temperatures of policy and action-selection, respectively. In the simulations below, we will simulate ‘one-shot’ experiments, in which there is no time-sensitive updating of precision, but we will illustrate the role of the hyperprior on precision (*β* and *α*) to simulate stochasticity or ‘random exploration’ in behaviour.

Using the same definition of free energy as in equation 1), but now with respect to the approximate posterior under a given policy, we obtain (Parr & Friston, 2018):

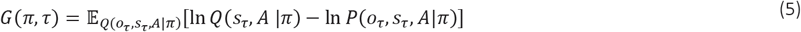

Defining *Q*(*o*_*τ*_, *s*_*τ*_, *A*|*π*) ≜ *P*(*o*_*τ*_|*s*_*τ*_, *A*)*Q*(*s*_*τ*_|*π*)*Q*(*A*) and applying the mean-field approximation, we obtain (Friston et al., 2017):

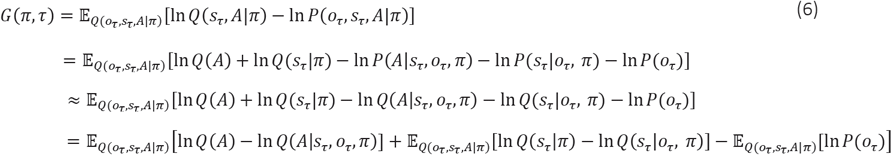

Finally, by applying the definition *Q*(*o*_*τ*_, *s*_*τ*_|*π*) = *Q*(*s*_*τ*_|*π*)*P*(*o*_*τ*_|*s*_*τ*_) again, we can define the value of a policy as:

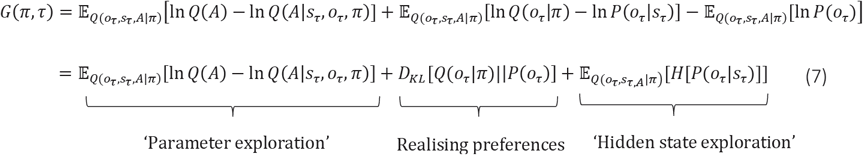

Here, *Q*(*o*_*τ*_, *s*_*τ*_, *A*|*π*) is the posterior predictive distribution over hidden states and their outcomes under a particular policy. Importantly, this formulation of behaviour predicts that choices will be governed by three principles; namely, minimising uncertainty about model parameters (*parameter exploration* or *novelty*), minimising uncertainty about hidden states (*hidden state exploration* or *salience*) and obtaining preferred outcomes (*realising preferences* or *goals*), which is defined as minimising the difference between predicted outcomes under a policy, *Q*(*o*_*τ*_|*π*), and preferred outcomes, *P*(*o*_*τ*_) (as defined in Figure 2C).

**Figure 2.**
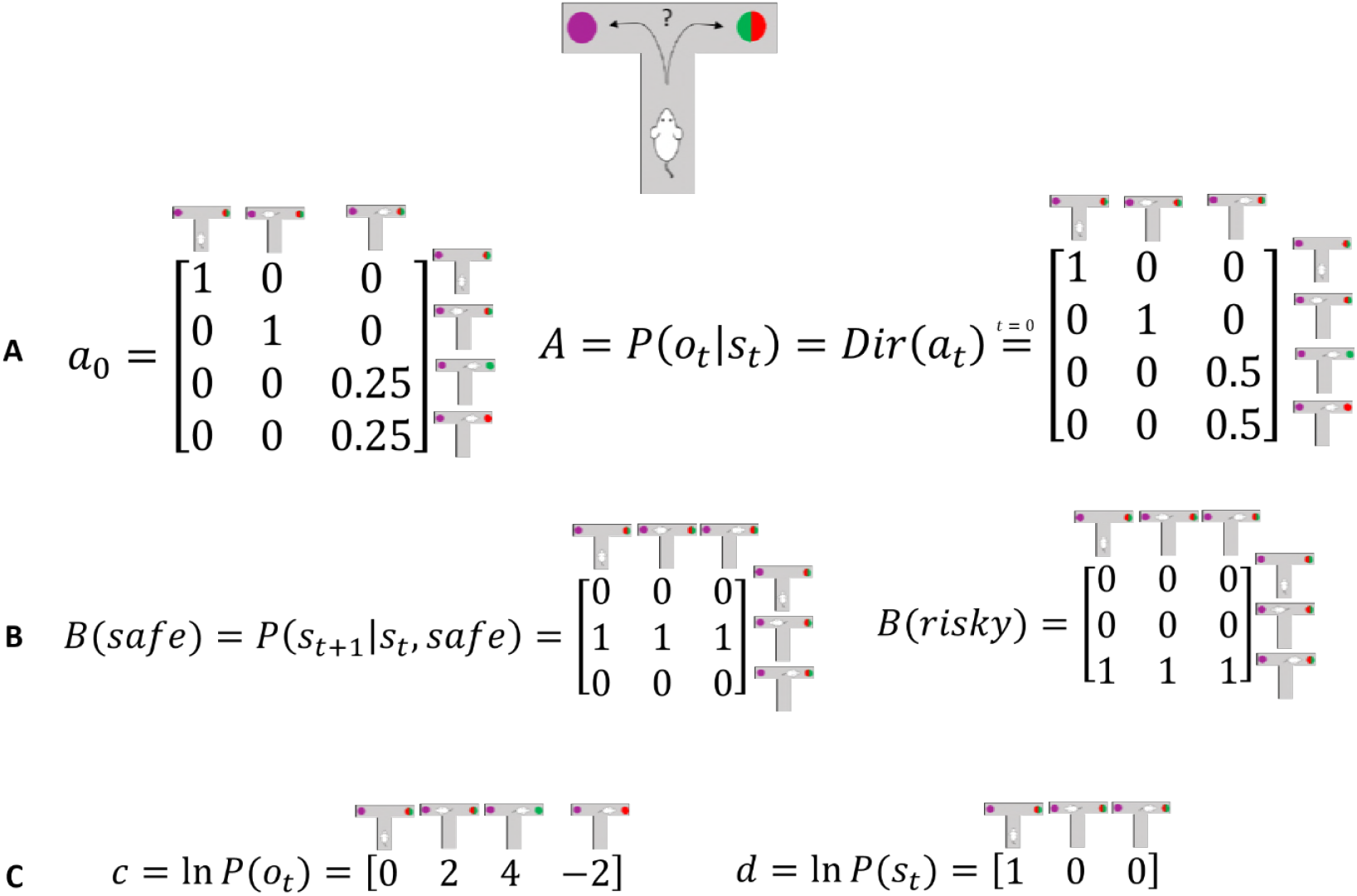
Generative Model of a T-maze task, in which an agent (e.g., a rat) has to choose between a safe option (left arm) and an ambiguous risky option (right arm). There are three different states in this task reflecting the rat’s location in the maze; namely, being located at the starting position or sampling the safe or risky arm. Further, there are four possible observations, namely being located at the starting position, obtaining a small reward in the safe option, obtaining a high reward in the risky option and obtaining no reward in the risky option. **A)** The A-matrix (*observation* or *emission* model) maps from hidden states (columns) to observable outcome states (rows, resulting in a 4×3 matrix). There is a deterministic mapping when the agent is in the starting position or samples the safe reward. When the agent samples the risky option, there is a probabilistic mapping to receiving a high reward or no reward. The A-matrix depends on concentration parameters *a* that are updated due to observing transitions between states and observations (in this example: receiving a high or no reward in the risky option), where *a*_0_ reflects the prior concentration parameters without having made any observation yet. **B)** The B-matrix encodes the transition probabilities, i.e. the mapping from the current hidden state (columns) to the next hidden state (rows) contingent on the action taken by the agent. Thus, we need as many B-matrices as there are different actions available to the agent (3 in this task: stay at the starting position, choose safe, choose risky). Here, the action simply changes the location of the agent. C) The c-vector specifies the preferences over outcome states. In this example, the agent prefers (expects) to end up in a reward state and dislikes to end up in a no reward state, whereas it is somewhat indifferent about the ‘intermediate’ states. Note that these preferences are (prior) beliefs or expectations are defined in log space, for example the agent beliefs that visiting the high reward state is exp(4) ≈ 55 times more likely than the starting point (exp(0) = 1) at the end of a trial. The d-vector specifies beliefs about the initial state of a trial. Here, the agent knows that its initial state is the starting point of the maze.

Note that the first term in the equation above reflects the mutual information between beliefs about model parameters before and after making an observation The notion of finding policies that maximise mutual information is equivalent to maximising (expected) Bayesian surprise (Itti & Baldi, 2009), where Bayesian surprise is the divergence between posterior and prior beliefs about hidden causes. Because mutual information cannot be less than zero, it disappears when the (predictive) posterior ceases to be informed by new observations. This means that ‘active learning’ will search out observations that resolve uncertainty about the world (e.g., foraging to resolve uncertainty about the reward probability of a risky option). However, when there is no posterior uncertainty – and the agent is confident about the structure of the world – there can be no further information gain and preferences over outcomes (i.e., rewards or utility) will dominate policy selection. This resolution of uncertainty is closely related to satisfying artificial curiosity (Schmidhuber, 1991; Still & Precup, 2012) and the ‘value of information’ (Howard, 1966). The third term of the value of a policy, on the other hand, penalises a policy for the expected entropy of the mapping between (hidden) states and observations. This term quantifies how well agents can infer the underlying cause of an observation, and motivates agents to seek observations with low ambiguity with respect to this mapping. Taken together, these two terms predict that (novel) policies will be preferred if they allow agents to optimise the parameterisation of their observation model and at the same time make (salient) observations that enable precise inference about the state of the world, given their observation model.

Actual updates of an agent’s observation model (A-matrix) at time-point *t* take place via updating concentration parameters with respect to current observations and an individual learning rate *η* (Friston et al., 2016, 2017):

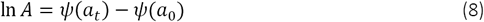

where *a*_*t*_ reflects the update of the concentration parameters depending on the observed state-outcome mapping at trial *t, a*_*t*_ = *a*_*t*-1_ + *η* · ∑_*t*_ *o*_*t*_⊗*s*_*t*_ (⊗ is the cross-product), and *a*_0_ reflects the (prior) concentration parameters at the beginning of the experiment, with Ψ referring to a psi- or digamma-function (i.e., a column-wise normalisation of concentration parameters). Note that *a* refer to the concentration parameters specifying an agent’s observation model via *P*(*A*) = *Dir*(*a*). Effectively, equation 8 implies that learning of the observation model takes place by counting the number of transitions from one particular hidden state to a particular outcome, modulated by an individual learning rate.

From the perspective of this paper, the key terms that define the value of a policy are the opportunities for information gain (i.e., novelty) pertaining to the mapping between hidden states and outcomes, and the expected entropy of the mapping from states to observations (i.e., salience). The former reflects an agent’s uncertainty about model parameters, whilst the latter reflects an agent’s uncertainty about hidden states. These two terms imply that policies will be preferred if they resolve uncertainty about the way in which hidden states generate outcomes (‘parameter exploration’) and about the hidden states underlying observations (‘hidden state exploration’). Interestingly, these two tendencies can make opposing predictions about behaviour. ‘Parameter exploration’ predicts that agents actively seek novel combinations of hidden states and outcomes, because they enable learning about the way in which outcomes are generated. ‘Hidden state exploration’ predicts that agents actively seek (known) salient observations that allow them to unambiguously infer the underlying hidden states. We will explore this dialectic between ‘active learning’ and ‘active inference’ in the following simulations.

## Model parameter exploration

In this section, we simulate behaviour with and without novelty seeking, curiosity-driven *active learning* that aims to acquire knowledge about the structure of a task (first term in equation 7). We will simulate a simple experiment, where an agent has to choose between a safe and a risky option, such as a rat in a T-shaped maze being forced to choose between the left and right arm to look for reward. We assume that the agent knows that it can only sample one of the two arms and that one arm (left in Figure 2) contains a certain small reward whereas the other arm (right in Figure 2) contains an uncertain, high reward. Importantly, however, the agent does not know about the reward probabilities in the uncertain arm in the beginning of the experiment, but can learn about these contingencies by updating its observation model via experience-dependent learning.

### Model Structure

To simulate behaviour, we need to specify the parameterisation of the model, which has been described in detail in previous work (Friston et al., 2016, 2017). In this task, we need to define a hyperprior on the precision of policy (choice) selection (*β* in Figure 1) and a prior on the precision of action selection (*α*). These parameters reflect the randomness of policy and action selection, respectively. Unless otherwise specified, we have set *β* to a (standard rate parameter) value of 1 and *α* to a value of 4. As shown in Figure 2, we define three different states in this task – as determined by the rat’s location in the maze; namely, being located at the starting position or sampling the safe or risky arm. Further, we define four possible observations; namely, being located at the starting position, obtaining a small reward in the safe option, obtaining a high reward in the risky option and obtaining no reward in the risky option. The ***A***-matrix (observation model) then determines the mapping from states to observations, while the ***B***-matrix (transition probabilities) specifies the mapping between hidden states given an action. Further, we need to specify an agent’s expectations over observations that reflect its preferences. These expectations are encoded in a **c**-vector, which we have set to *c* = [0 2 4 −2] in the following simulations, reflecting an agent’s preference for being in the starting position, obtaining a safe reward, obtaining a high reward and obtaining no reward in a risky option, respectively. These preferences are defined as the agent’s log-expectations over outcomes. For example, the definition of these preferences implies that the agent beliefs that visiting the high reward state is **exp(4) ≈ 55** times more likely than visiting the starting point **(exp(0) = 1)** at the end of a trial. The **d**-vector encodes an agent’s expectations about the initial state, which was defined to reflect full certainty about starting each trial in the starting position of the maze. In simulations that include learning, we set the initial concentration parameters for obtaining a high reward (or not) to 1/4 (i.e., position (3,3) and (4,3) in the A-matrix in Figure 2), and these concentration parameters are updated according to a learning rate *η*, which was set to 0.5. Figure 2 illustrates the architecture of the generative model of this task.

### Active Learning

Figure 3 illustrates an experiment that was simulated under active learning with an underlying high-reward probability of 50%. The bottom panel illustrates the evolution of beliefs (concentration parameters) about the underlying emission probabilities of the task for every trial of the experiment, which in turn determine policy selection as illustrated in the first panel. Note that at the start of the experiment, the agent assigns equal probability to receiving a high reward and no reward at the risky option, but these beliefs have very low certainty (i.e., very small concentration parameters, see mapping to ‘high reward’ and ‘low reward’ in risky option at the ‘start’ position in third panel of Figure 3). This leads the agent to mainly explore and gather information in the beginning of the experiment by choosing the risky option. Interestingly, after trial ten, the agent (correctly) assigns a probability of 50% to a high reward in the risky option, but now with higher confidence (i.e., larger concentration parameters). Consequently, the agent now prefers to exploit and sample the safe option, driven by both the expected value of this option and a preference for visiting unambiguous states.

**Figure 3.**
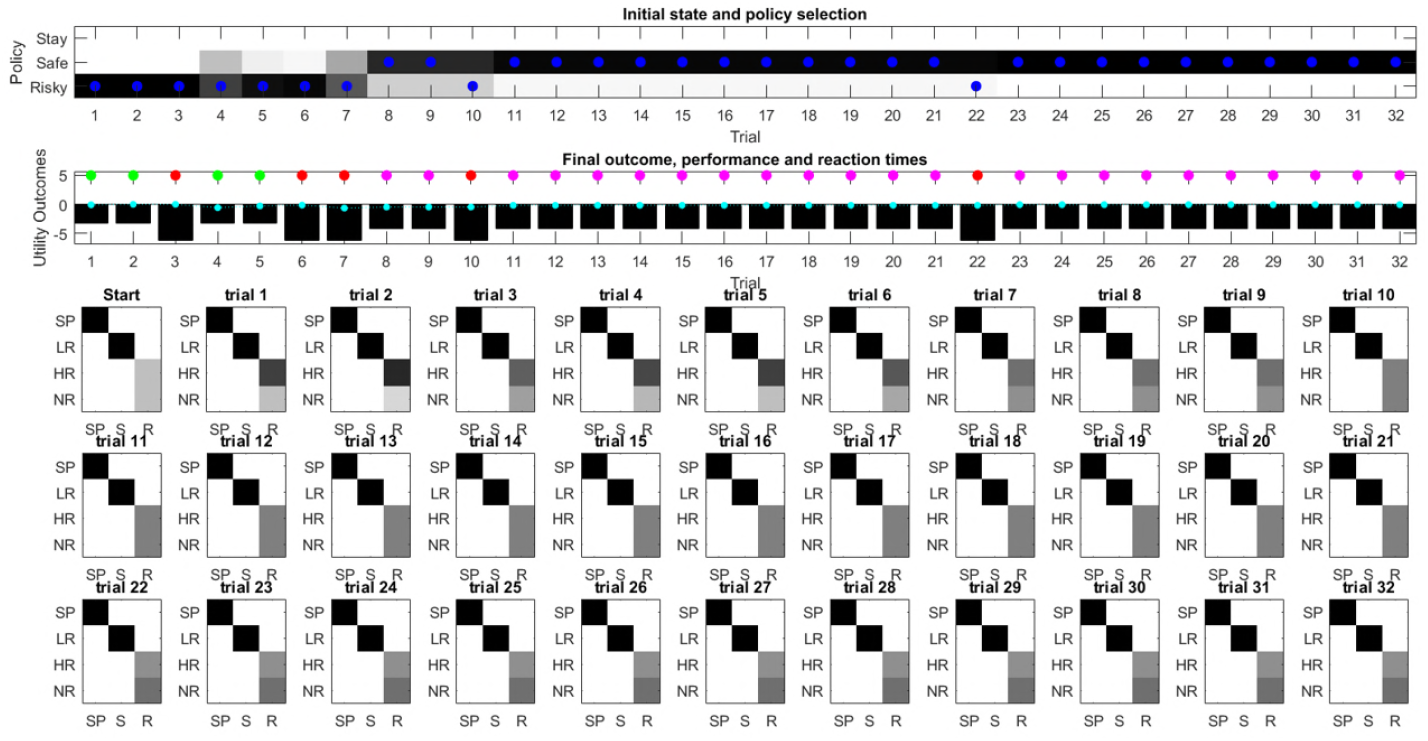
Simulated responses during learning: This figure illustrates responses and belief updates during a simulated experiment with 32 trials. The first panel illustrates whether the agent sampled the safe or risky option as indicated by the blue dots, as well as the agent’s beliefs about which action to select. Darker background implies higher certainty about selecting a particular action. The second panel illustrates the outcomes at each trial, the utility of each outcome and simulated choice conflict. Outcomes are represented as coloured dots, where magenta refers to a small and safe reward, green to a high reward and red to no reward in the risky option. Cyan dots reflect the simulated choice conflict defined as the entropy of beliefs over choices (i.e., the columns in the panel above). Black bars reflect the utilities of the outcome. Note that these utilities are defined as log-expectations over outcomes (see main text and Figure 2), thus a value closer to zero reflects higher utility of an outcome. Panels three to five illustrate the updates of the concentration parameters of the observation model, which specify the mapping from hidden states (columns: ‘SP’ = starting point, ‘S’ = safe option, ‘R’ = risky option) to observed outcomes (rows: ‘SP’ = starting point, ‘LR’ = low reward, ‘HR’ = high reward, ‘NR’ = no reward). In this example, the simulated agent makes predominantly curious and novelty-seeking choices in the beginning of the experiment. After the tenth trial, the agent is confident that the risky option provides a probability of 0.5 for receiving a high reward, which compels her to choose the safe option afterwards.

Figure 4 illustrates the same task but with a reward probability of 0.75 for the risky option. Here, after a similar number of exploration trials as in Figure 3, the agent becomes confident that it should select the risky option, given its higher expected value. This can be seen by the fact that the agent continues to select the risky option (blue dots) with high confidence (shaded area behind blue dots, first panel) because the risky option is mostly rewarded (green dots, second panel).

**Figure 4.**
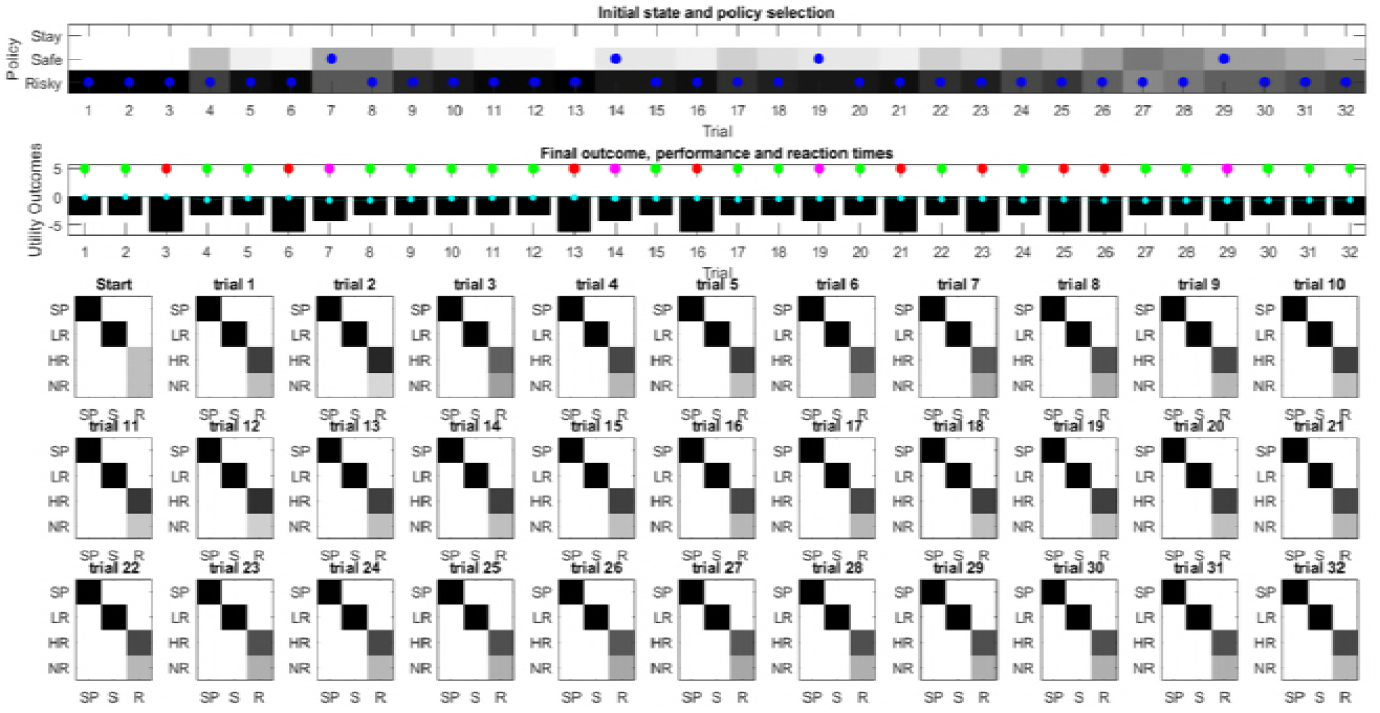
Learning a high reward probability in the risky option. Same setup as in Figure 3, but now the true reward probability of the risky option is set to 0.75. This means that after sampling the risky option in the beginning of the experiment and learning about the high reward probability of that option (as shown in panel three), the agent becomes increasingly certain that the risky option has a high probability of a reward, i.e., the mapping between the state ‘risky option’ to the observation ‘high reward’ becomes stronger (panels three to five). This compels the agent to continue sampling the risky option and only rarely visiting the safe option with low certainty, as illustrated in panel one.

### The role of precision in active learning

The above simulations highlight an important aspect of exploratory behaviour, namely behaviour that is goal-directed and aims at reducing uncertainty about a specific part of an agent’s model, in this example the part of the A-matrix (i.e., the observation or emission function) that specifies the mapping from sampling the risky option to obtaining a high or low reward. This means that the agent tries to gain insight into a particular part of the structure of world that it is unsure about. Importantly, this predicts that this sort of exploratory behaviour will be most prevalent if there is high uncertainty about the structure of a task, such as in the beginning of a game (cf., Figure 7). This also suggests an important confound when investigating the influence of reward and uncertainty on behaviour; namely, the fact that the rewarding options will often be associated with the lowest uncertainty because they are sampled most frequently (Wilson, Geana, White, Ludvig, & Cohen, 2014b), which highlights the importance of analysing behaviour at the beginning of an experiment when there is high uncertainty about all available options (Gershman, 2018b, 2018a).

As illustrated earlier, goal-directed information-gain can be contrasted with random exploration, such as in simple ε-greedy or softmax choice rules where the degree of randomness is governed by an inverse temperature parameter (Sutton & Barto, 1998). In its simplest form, random exploration implies that exploratory behaviour will not be sensitive to an agent’s uncertainty about different options or its uncertainty about different parts of the world. This implies that such behaviour will not decrease uncertainty per se but may cause ‘accidental’ belief-updating due to random or stochastic selection of different policies. Here, this sort of behaviour is controlled by the precision of policy and action selection (see equation 4). This means that random exploration can be understood as imprecise behaviour. Importantly, the precision of behaviour does not depend on an agent’s uncertainty about the world, such that there is no predicted relationship between ‘random exploration’ and the time-course of an experiment (see below). Figure 5 illustrates the effects of highly imprecise (*β* = 2^3^, Figure 5A) and highly precise (*β* = 2^-3^, Figure5B) types of behaviour. Note that the expected value of precision is the inverse of *β*, i.e., 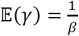 (Figure 1).

**Figure 5.**
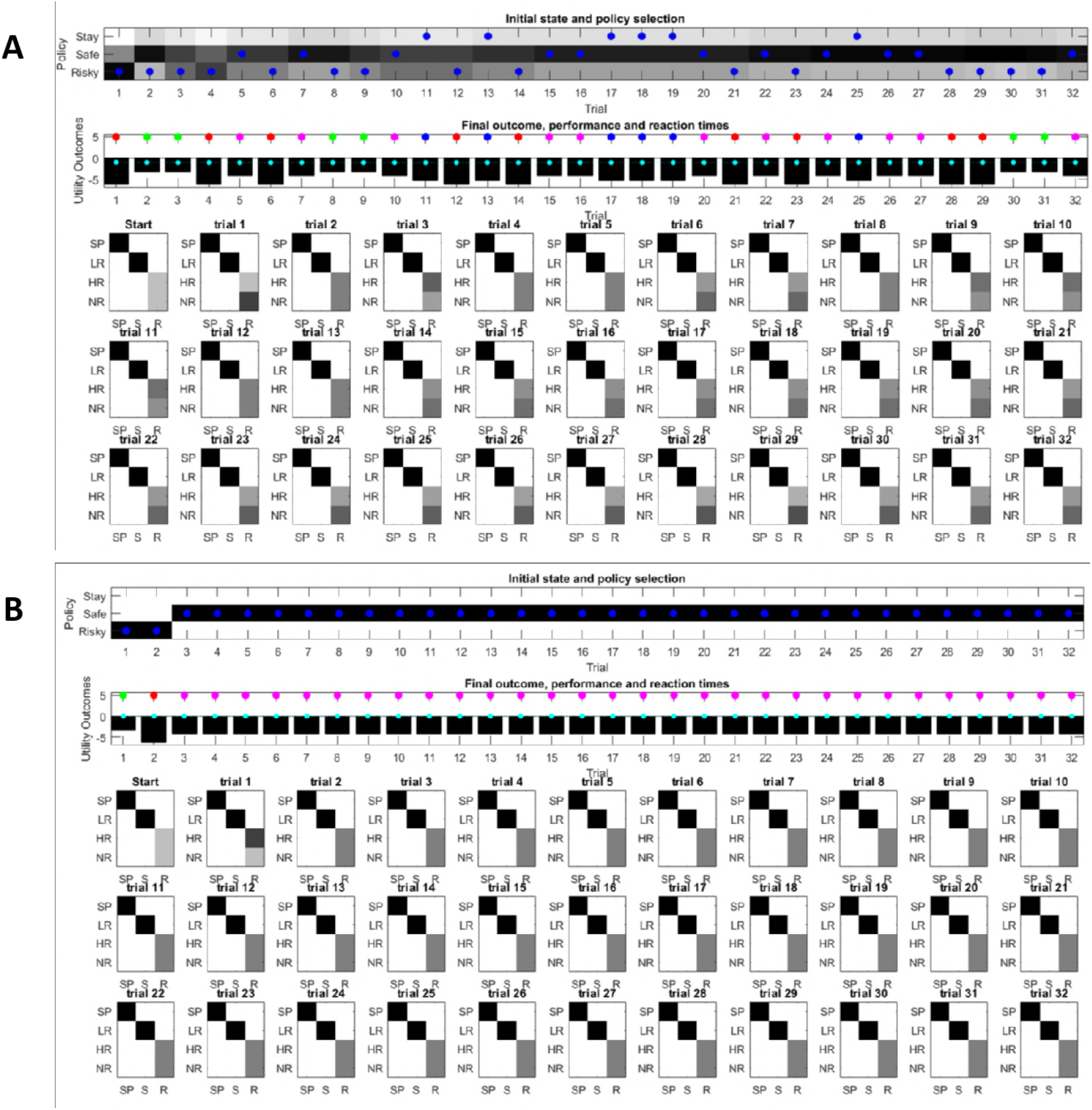
Effects of precision on behaviour. Same setup as in Figure 3, but now with varying levels of precision. A) A high degree of random exploration results from very imprecise behaviour (*β* = 2^3^), whereas B) highly precise behaviour (*β* = 2^-3^) results in very low randomness in behaviour.

### Broken ‘parameter exploration’

Equation 7 shows that the ability to learn about the environment and minimise uncertainty is a determining factor of the value of policies. This can be illustrated by disabling any influence of such learning on policy evaluation, as shown in Figure 6. In this case, policies cannot be distinguished in terms of their uncertainty reduction about model parameters. Consequently, the only factors that determine the value of policies are visiting preferred and unambiguous outcomes. This means that agents will not exhibit active learning, and the only way to learn about the environment is by accidently (randomly) sampling a non-preferred option. Figure 6 illustrates this problem: here, the true reward probability of the risky option is 0.75, but in the absence of any active learning, the agent can only find out about the value of the risky option by randomly sampling this alternative. Thus, if the agent shows very precise (non-random) behaviour (Figure 6A), it is very unlikely to discover that the risky option is better than the safe option, and only by showing very imprecise behaviour (Figure 6B) the agent will be able to develop a (weak) preference for the risky option.

**Figure 6.**
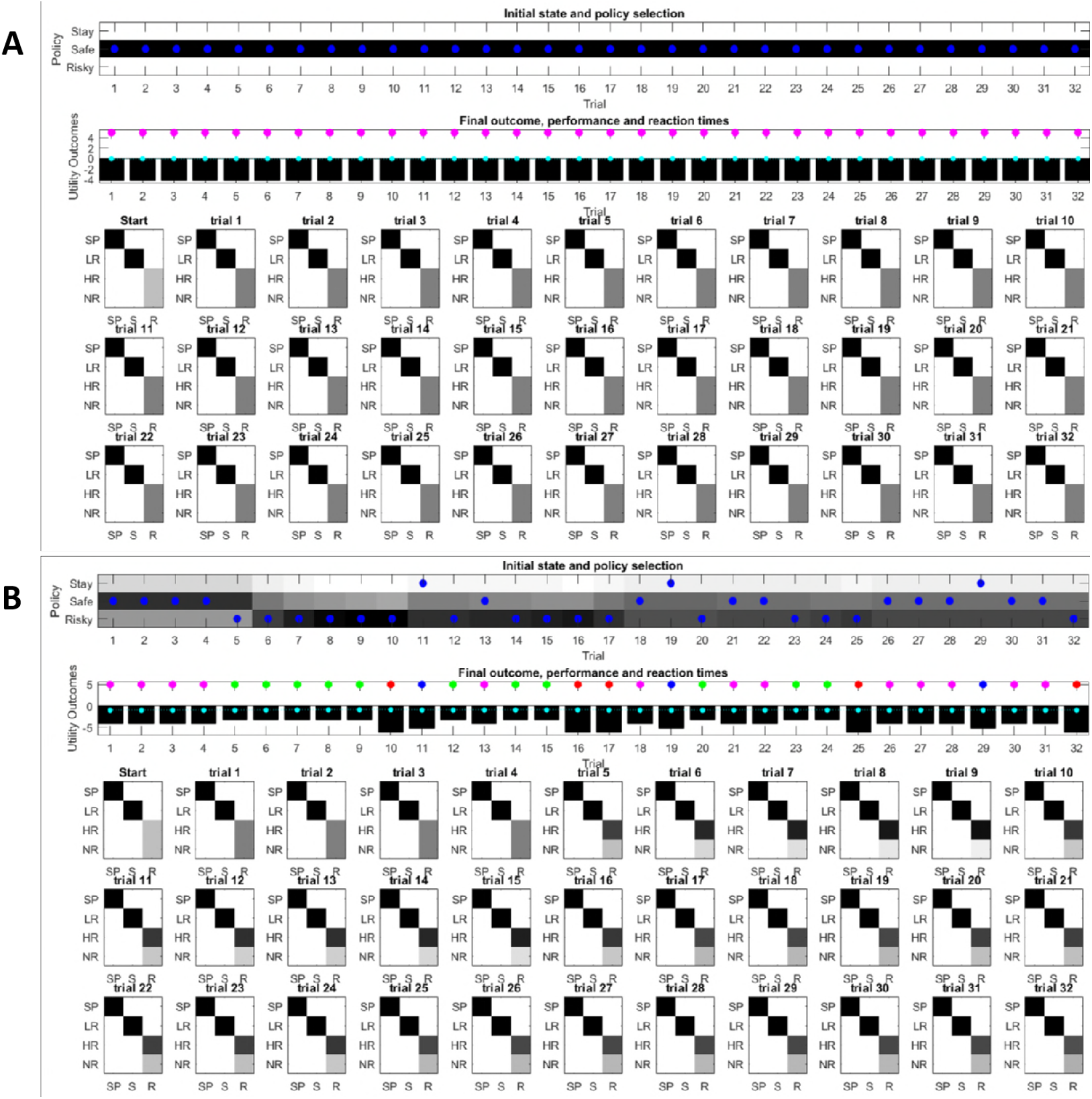
‘Broken’ parameter exploration. Same setup as in Figure 3, but now with a true reward probability of 0.75 and no active learning as a determinant of the value of policies (first term of equation 7). A) If behaviour is very precise (*β* = 2^-3^), the agent will never find out that the risky option is more preferable than the safe option, because there is no active sampling of its environment. B) In contrast, if the agent’s behaviour has a higher degree of randomness (low precision, *β* = 2^3^), then it will eventually learn about the reward statistics in the risky option from randomly sampling this alternative, and infer that it is preferable over the safe option.

### Time courses of exploratory behaviour

A general problem when investigating the role of exploration in value-based decision-making is that if an agent is allowed to move around freely, there will be a relationship between the reward statistics of an option and its associated uncertainty. Rewarding arms will be associated with a lower level of uncertainty simply because they are sampled more often (Gershman, 2018b, 2018a; Wilson et al., 2014). To compare different computational architectures that might underlie exploratory behaviour and information-gain, it is therefore important to investigate the time-course of behaviour, as illustrated in Figure 7 based on a true reward probability of 0.5 in 1000 simulations of the task described in the previous figures. Figure 7A illustrates the time-course of behaviour under active learning conditioned on the concentration parameters of the A-matrix (observation model, left panel) and conditioned on the trial-number (right panel).

**Figure 7.**
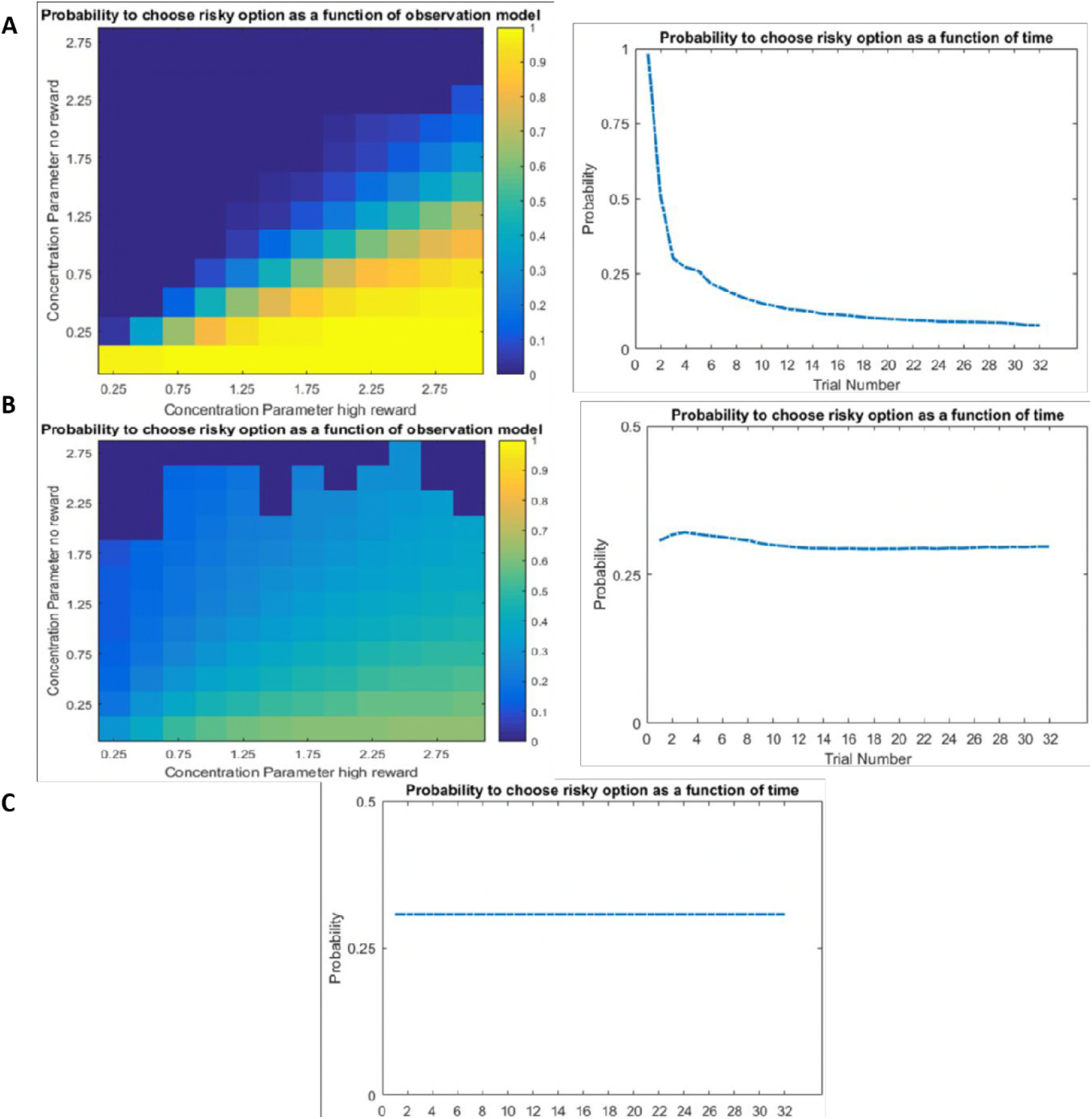
Time-course of active learning and random exploration. Simulations of 1000 experiments with 32 trials each under a true reward probability of 0.5. A) In active learning, the probability to choose the risky (uncertain) option is high if there is high uncertainty about this option (left panel, probability to choose risky option as a function of the concentration parameters for high reward and no reward in the risky option) at the beginning of a task (right panel, average probability to choose the risky option as a function of time). Note how the probability of choosing the risky option decreases as the agent becomes more certain that the true reward probability of the risky option is 0.5 (diagonal of left panel). B) When there is no active learning but high randomness (low prior precision, *β* = 2^3^), there is no preference for uncertainty-reduction at early trials, and the probability to choose the risky option quickly converges as the estimate of the true reward probability converges to 0.5 due to random sampling of the risky option. C) In the absence of any learning, the probability to choose the risky option is constant and reflects the precision or randomness in an agent’s generative model. Note the smaller scale of the y-axis in B) and C) compared to A) for better readability.

Unsurprisingly, we observe that the agent strongly prefers to choose the risky option when she believes that the reward probability is high (right bottom corner of left panel in Figure 7A) and strongly prefers to choose the safe option if the probability of a high reward is low (left upper corner in left panel in Figure 7A). More interestingly, we also observe a gradient across the diagonal, such that agents have a strong preference to choose the risky option if there is high uncertainty about its reward contingencies (i.e., both concentration parameters of the ***A***-matrix are low, lower left corner of left panel in Figure 7A). In contrast, the probability to choose the risky option is very low if the agent is very certain that the probability to receive a high reward is 0.5 (i.e., both concentration parameters of the ***A***-matrix are high, upper right corner of left panel in Figure 7A). In line with this, the probability of choosing the risky option over time under active learning shows that there is a very high preference for sampling the risky (uncertain) option in the beginning of a trial, which then monotonically decreases over time (right panel in Figure 7A).

Figure 7B illustrates the time-course of behaviour without active learning but with a high degree of random exploration (low prior precision), where the only way to learn about the true reward probabilities is by randomly sampling the risky option. The pattern of the left panel of Figure 7B looks like a noisier version of the left panel of Figure 7A. Aside from the larger randomness in behaviour, there is also an important difference when uncertainty about the true reward statistics is high (lower left corner): in the absence of active learning, there is no preference for the risky option when the relevant concentration parameters of the A-matrix are both low (lower left corner in left panel of Figure 7B). This also becomes apparent when looking at the time course of choosing the risky option, such that there is no initial preference for the risky option reflecting uncertainty reduction in the beginning of a trial. Rather, the probability to select the risky option remains relatively stable across trials and reflects the overall level of randomness in behaviour.

Finally, Figure 7C illustrates the time course of the probability to choose the risky option if there is no learning at all (i.e., the concentration parameters of the A-matrix do not change). In this case, the probability to choose the risky option is constant and simply reflects the precision of individual behaviour.

## Hidden state exploration

In this section, we illustrate a second type of behaviour that aims at gaining information about the world, namely exploring about hidden states of a task, as reflected by the third, salience term of equation 7. In contrast to ‘model parameter exploration’, which motivates *active learning* to reduce uncertainty about an agent’s model of the world, ‘hidden state exploration’ motivates *active inference* to form accurate beliefs about the current state of the world, based on an agent’s model of the task. One example of this behaviour is inferring the current context, which we illustrate in the following simulations, using a slightly adjusted version of the previous task. We now assume that the agent has learned that she could be in two possible (hidden) states in this task, namely either in a context where the risky option provides high or low probability for obtaining a reward, but this contextual information is hidden from her. However, in this version of the task, she can also choose to sample a cue before choosing the safe or risky option, which tells her about the reward probabilities (i.e., context) of the current trial.

### Model Structure

The generative model of the ‘hidden state exploration’ task is illustrated in Figure 8. We have used the same formalisation and parameter settings (with *β* = 1 and *α* = 16 unless otherwise specified) as in the previous model, except that the agent now performs inference about sampling the safe or risky option directly, or sampling a cue first that signifies the current context, namely a high (75%) or a low (25%) probability to obtain a reward in the risky option. In comparison to the previous generative model illustrated in Figure 2, this increases the size of the state space by the additional cue location and the (hidden) context factor, resulting in eight different (hidden) states (columns of A-matrix in Figure 8). The B-matrix encodes the transitions between different locations from the starting position of the maze; namely, sampling the cue, the safe option, or the risky option. The c- and d-vectors are defined analogously to the previous example, except that the d-vector now reflects a uniform prior about starting the maze in one of the two contexts. We did not include any curiosity-driven learning in these simulations, except that we allowed for experience-based updates of the d-vector in two simulations (Figure 10 and 11), which describe a task in which the true state of the task can be learned gradually. Updates of the (concentration parameters of the) d-vector are implemented analogously to the updates of the A-matrix in the ‘parameter exploration’ example above. Note that, in principle, such updates would also allow the agent to continuously learn about the current reward probabilities of the risky option without sampling the cue first, analogously to the ‘model parameter exploration’ example. Importantly, however, parameter exploration will not work if the context changes rapidly, such as on a trial-by-trial basis. This provides an important illustration of the different time-courses of inference and learning, which we discuss in more detail below.

**Figure 8.**
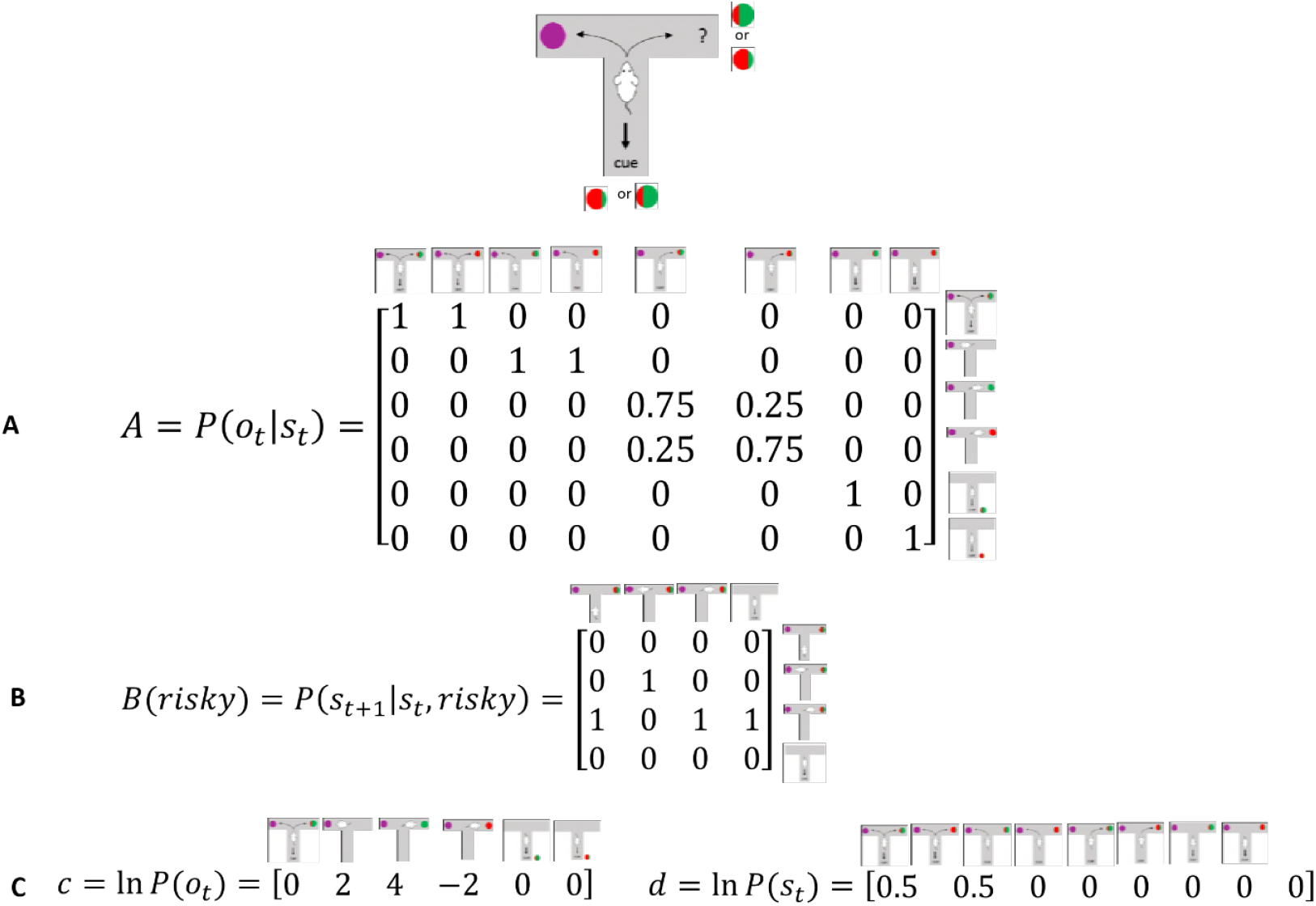
Generative Model of a T-maze task, in which an agent (e.g., a rat) has to choose between a safe option (left arm) and a risky option (right arm). In contrast to the previous task, the rat can now be in two different contexts that define the reward probability of the risky option, which can be high (75%) or low (25%). Besides sampling the safe or risky option, it can now also sample a cue that signifies the current context. This results in a state space of eight possible states, defined by the factors location (starting point, cue location, safe option, risky option) and context (high or low reward probability in risky option). Further, there are seven possible observations the agent could make, namely being at the starting position, sampling the safe option, obtaining a/no reward in the risky option, and sampling the cue that indicates a high/low reward probability. **A)** The A-matrix (*observation* or *emission* model) maps from hidden states (columns) to observable outcome states (rows, resulting in an 8×7 matrix). There is a deterministic mapping when the agent is in the starting position, samples the safe reward or samples the cue. When the agent samples the risky option, there is a probabilistic mapping to receiving a high reward or no reward that depends on the current context. In contrast to the previous example, no updates of the A-matrix take place in this task. **B)** The B-matrix encodes the transition probabilities, i.e. the mapping from the current hidden state (columns) to the next hidden state (rows) contingent on the action taken by the agent, which simply changes the location of the agent. For simplicity, only the transition probabilities for the factor location are shown, which replicate across the two contexts (resulting in an 8 × 8 transition matrix). **C)** The c-vector specifies the preferences over outcome states. In this example, the agent prefers (expects) to end up in a reward state and dislikes to end up in a no reward state, whereas it is somewhat indifferent about the ‘intermediate’ states (starting position or cue location). The d-vector specifies beliefs about the initial state of a trial. Here, the agent knows that its initial state is the starting point of the maze, but has a uniform prior over the two contexts. In experiments where the context is stable, this uniform prior can be updated to reflect experience-dependent expectations about the current context.

### Active Inference

Figure 9 illustrates ‘hidden state exploration’ in an experiment, where the current context cannot be learned, i.e. changes randomly on a trial by trial basis. In that case *active inference* predicts that the agent will always sample the cue at the beginning of every trial to reduce ambiguity about the current hidden state (context) (first and third panel of Figure 9). The subsequent behaviour in a trial depends on the information obtained at the cue. If the cue signifies a context with high reward probability (dark green dots in second panel of Figure 9), the agent will choose the risky option. In contrast, if the cue indicates a context with a small reward probability, she will choose the safe option.

**Figure 9.**
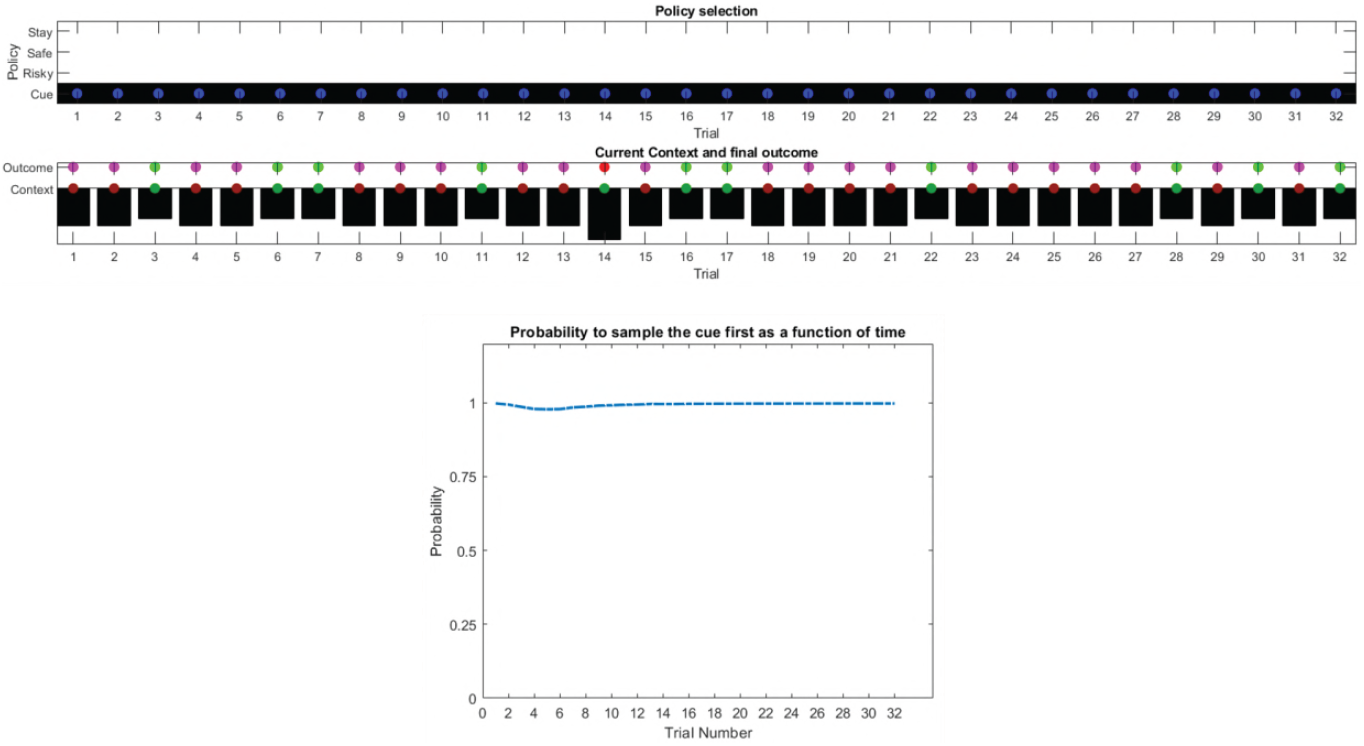
Simulated responses during inference: This figure illustrates responses during ‘hidden state exploration’ in a simulated experiment with 32 trials (first two panels) and based on simulations of 1000 experiments with 32 trials each (third panel). In this experiment, the current context, reflecting either high (75%) or low (25%) reward probability in the risky option, changes on a trial-by-trial basis and thus cannot be learned. The only way for the agent to gain information about the current context of a trial is by sampling a cue, which signifies the current context. As above, the first panel illustrates the choice of the agent at the beginning of a trial and the agent’s beliefs about action selection (darker means more likely). Note that the agent always chooses to sample the cue first before choosing the safe or risky option. The second panel illustrates the outcomes of every trial (magenta = safe option, green = high reward in risky option, red = no reward in risky option) and their utilities (black bars, closer to zero indicates higher utility). Note that a green or red outcome indicates that the agent has chosen the risky option after sampling the cue. Dark red and green dots indicate the current context as signified by the cue (dark red = low reward probability in risky option, dark green = high reward probability in risky option). Note that the agent only samples the risky option if the cue indicates a high reward context. Third panel illustrates the probability to sample the cue at the beginning of a trial, simulated for 1000 experiments with 32 trials each. There is a nearly 100% probability to sample the cue first at every trial, because in these simulations the context changed on a trial-by-trial basis.

This simulation illustrates an important difference to the active learning simulations above: in these simulations, there is nothing to be learned about the state of the world, because the current state changes randomly on a trial by trial basis. Thus, this task could not be solved by learning the reward-mapping of the risky option, because there is no knowledge about the reward statistics that could be carried over from one trial to the next. This highlights the necessity to perform trial-by-trial inference about the current state of the world, as opposed to continuous parameter learning.

Figure 10 illustrates simulations of the same task, but now with a stable context of a high reward probability in the risky option, allowing for experience-dependent updates of the agent’s prior over initial contexts (in the d-vector, cf. Figure 8) based on information obtained from the cue. In the first third of the experiment, we observe the same choice bias as in Figure 9, namely a preference to sample the cue first before choosing the safe or risky option. In this experiment, however, the agent always obtains the same information from the cue location, indicating a stable environment with a high reward probability in the risky option. Once the agent becomes confident enough in its beliefs about the current context, it starts to sample the risky option without sampling the cue first. Note that in contrast to the ‘parameter exploration’ simulations above, the agent updates its beliefs based on the (hidden state) information provided by the cue, not the actual outcome (i.e., obtaining a reward). This can be seen in the belief-updating after trial one, for instance: the agent samples the cue, which indicates a high reward probability context, and obtains no reward from the risky option. Despite the negative outcome, it increases its belief about being in the high reward context (third panel of Figure 10, columns one and two), due to the information obtained from the cue.

**Figure 10.**
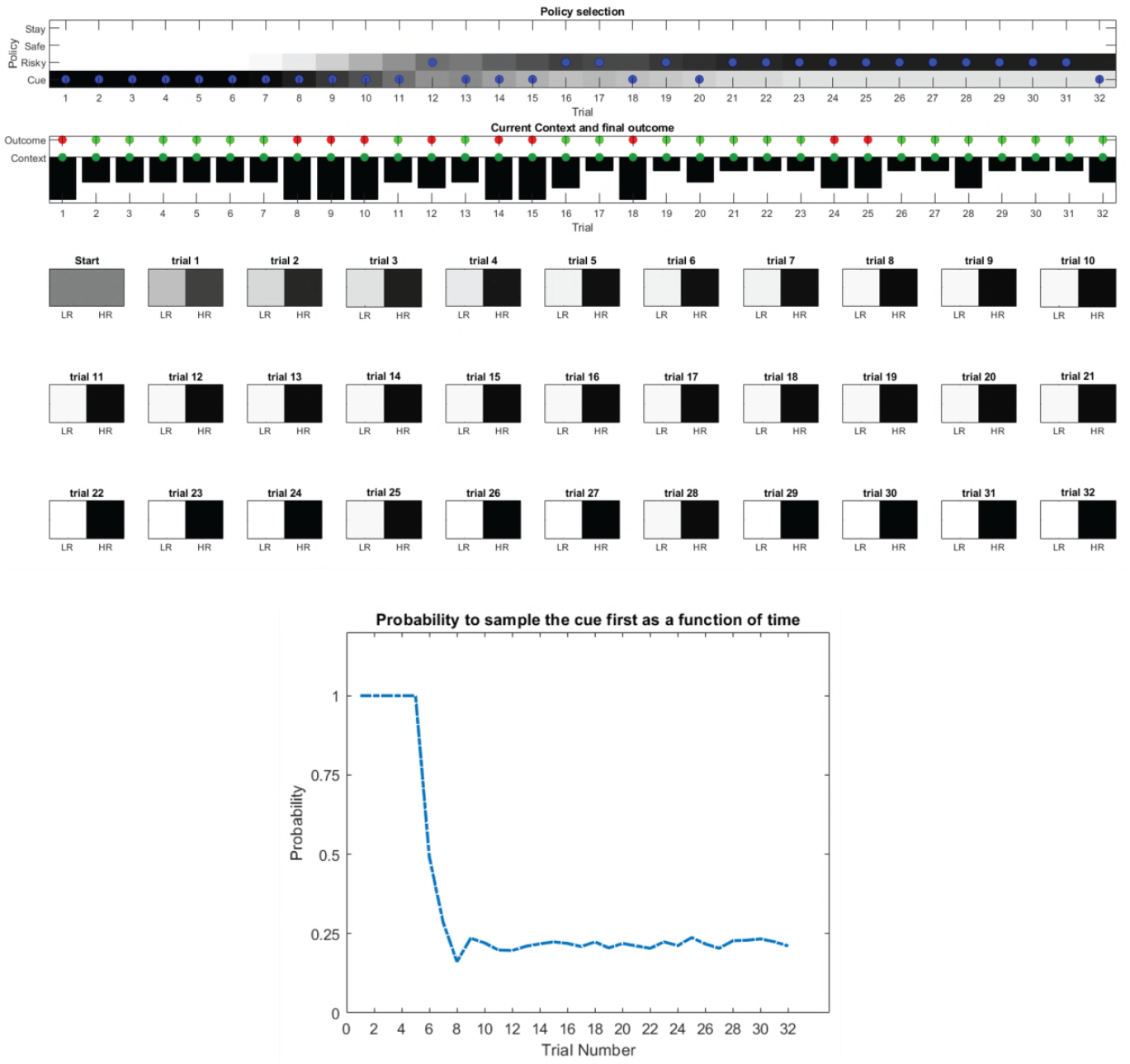
Simulated responses during inference in a stable context: Same setup as in Figure 9, but now with a constant context that indicates a high reward probability in the risky option. Belief updates (in the d-vector, cf. Figure 8) are illustrated in panels three to five (LR = low reward context, HR = high reward context, darker means more likely). Here, the agent becomes increasingly confident that it is in a high reward context, which compels it to sample the risky option directly after about one third of the experiment, whilst gathering information in the cue location in the first third of the experiment. The lowest panel illustrates the time-course of the probability to sample the cue first as a function of trial number in an experiment (in 1000 simulated experiments). The probability to sample the cue shows a sharp decrease once the agent has gathered enough information about the current context.

### Broken ‘hidden state exploration’

What happens if an agent fails to perform ‘hidden state exploration’? Figure 11 shows simulations of behaviour when information-gain about the hidden state is not considered during policy selection. This implies that the cue location has no informative value, and is equally preferable to the starting location of the maze (because they have the same utility, cf. c-vector of Figure 8). Analogously to behaviour illustrated in Figure 6, this implies that agents can only learn about a stable current context through random behaviour (illustrated herewith a decreased precision of action selection with *α* = 4), resulting in a constant low probability to sample the cue location across time, which reflects the overall level of randomness in an agent’s behaviour. In the example illustrated in Figure 11, the agent fails to acknowledge that there is a high reward probability in the risky option and continues to prefer the safe option.

**Figure 11.**
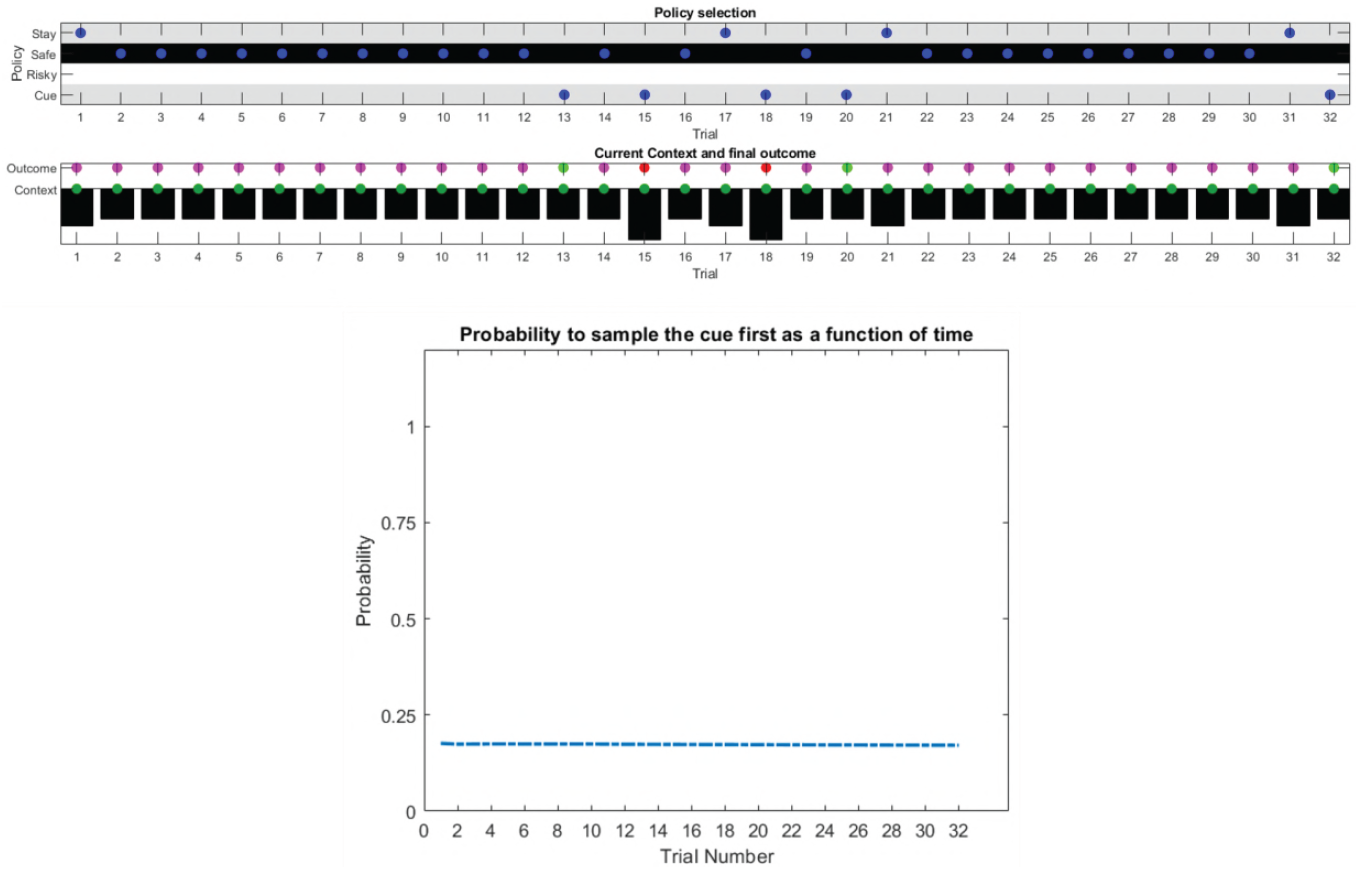
‘Broken’ hidden state exploration: Same setup as in Figure 10, but now without a ‘hidden state exploration’ bias in policy selection (third term of equation 7). The agent fails to learn that there is a constant high reward probability for the risky option because it does not gain information about the current hidden state (context). Consequently, it continues to prefer the safe option. The probabilities to sample different options (first panel) now simply reflect the agent’s prior preferences as encoded in the c-vector (cf., Figure 8). The time-course of the probability to sample the cue location (lower panel) now simply reflects the agent’s general level of randomness in behaviour.

## Comparing model parameter and hidden state exploration

In this final section, we provide direct comparisons of parameter exploration (*active learning*) and hidden state exploration (*active inference*) in different variants of the tasks introduced above. In these simulations, we use an identical parameterisation for these two types of behaviour (particularly with *α* = 8, *β* = 1) except that ‘parameter exploration’ (active learning) is only governed by the first two terms of equation 7, namely model updating and realising preferences based the generative model defined in Figure 2, and ‘hidden state exploration’ (active inference) is only governed by the last two terms of equation 7, namely realising preferences and minimising ambiguity based on the generative model defined in Figure 8. Further, we will contrast these types of goal-directed exploration to a ‘random exploration’ agent with a higher degree of stochasticity in its behaviour (*α* = 1), but no bias for (goal-directed) parameter or hidden state exploration, which will serve as a baseline for the other two types of exploratory behaviour. Thus, this agent will be solely governed by the (second) realising preferences term in equation 7, but can still update its model of the task due to randomly sampling different options. We will compare these agents in situations where the risky option is either advantageous (reward probability of 85%) or disadvantageous (reward probability of 15%). We use the average cumulative reward in 100 simulated experiments with 32 trials each as a measure of performance for these three agents, where we define a low reward as one food pellet and a high reward as four food pellets that could be obtained by the rat.

Figure 12 shows the behaviour of these three agents in the task illustrated at the top of Figure 2, where a rat has to choose between a certain safe and an uncertain risky option. In line with the previous simulations, we observe that the ‘parameter exploration’ agent quickly learns to prefer the risky option if there is a high reward probability (left upper panel of Figure 12) and to avoid the risky option if there is a low reward probability (right upper panel). The ‘random exploration’ agent also converges on these estimates, but much slower. Interestingly, we observe that the ‘hidden state exploration’ agent fails to adjust to the reward statistics of this task. This is because, from the perspective of this agent, there is no hidden state to explore that could be informative about the current reward statistics. The only way to learn about the statistics of the task would be by sampling observations that are a priori associated with high ambiguity. Such observations, however, are aversive for a pure *active inference* agent, because they are associated with a high entropy (ambiguity) in their mapping to underlying hidden states, which an *active inference* agent is compelled to *minimise*. Consequently, it will always sample the safe option in this task. This induces a performance pattern in which the ‘parameter exploration’ agent is superior to the other two agents if the reward probability in the risky option is high, but a similar performance level if the reward probability is low and the best course of action is to sample the safe option (left and right lower panel of Figure 12).

**Figure 12.**
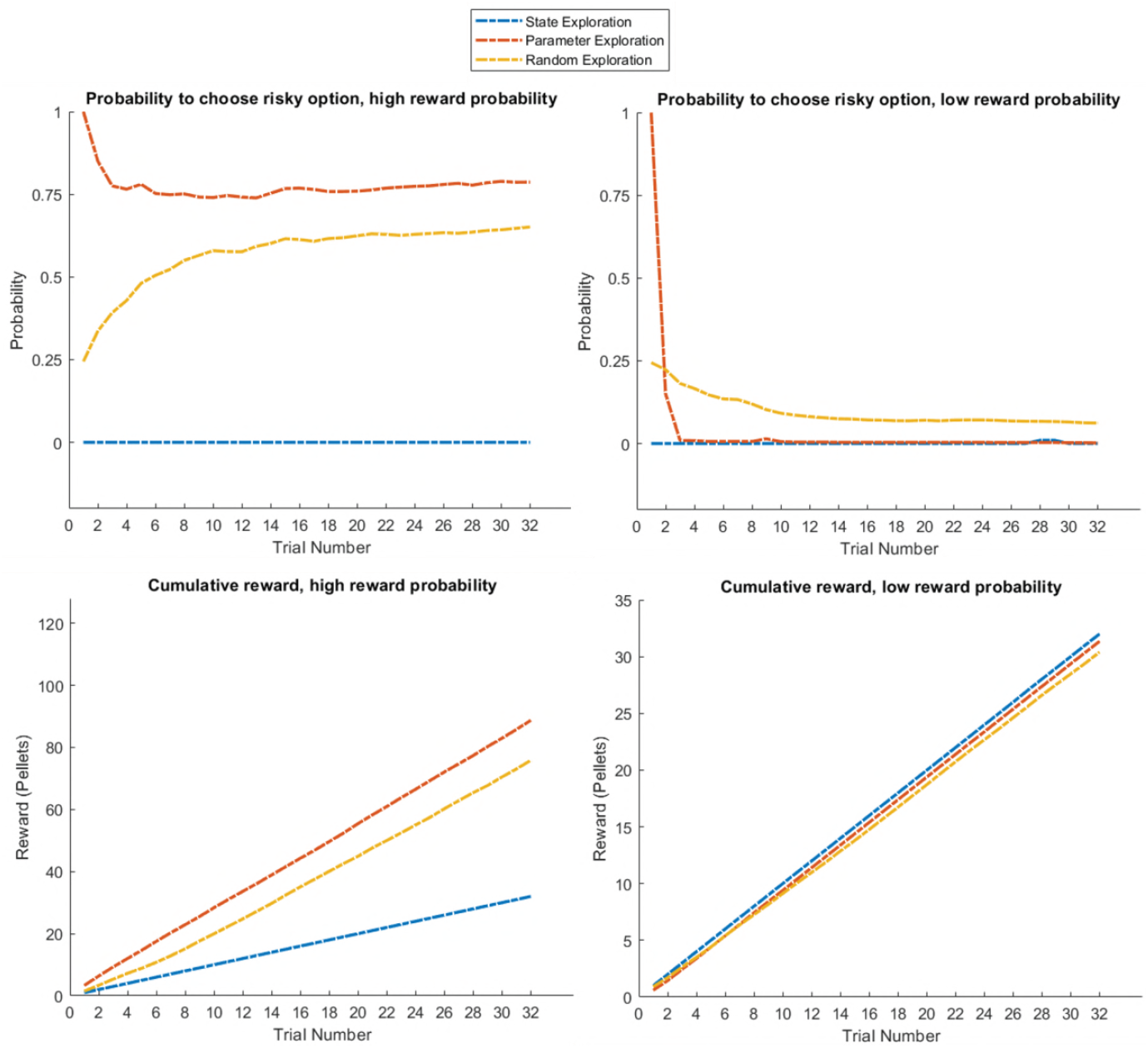
Response profiles of a ‘state exploration’, ‘parameter exploration’ and ‘random exploration’ agent in a task that requires learning. In the task described at the top of Figure 2, only the ‘parameter exploration’ agent flexibly adapts to the current reward statistics, whilst the ‘state exploration’ agent fails to form a representation of the task statistics. This is because optimal behaviour requires agents to learn about the task by sampling novel observations that are associated with high a priori uncertainty, which is the essence of *active learning* (parameter exploration) but not *active inference* (state exploration). Upper panel: probability for each of the three agents to choose the risky option if it is associated with a high (left, 85%) or low (right, 15%) reward probability. Lower panel: average cumulative reward (measured in pellets, where low reward = one pellet and high reward = four pellets) in 100 simulated experiments in a high (left) and low (right) reward probability setting, indicating an advantage for the ‘parameter exploration’ agent when the risky option is associated with a high reward probability.

Figure 13 compares the three agents in the task introduced in Figure 9, where the current context (high or low reward probability in the risky option) changes unpredictably on a trial-by-trial basis, but can be inferred from sampling a cue that signifies the current context. This illustrates the opposite situation to Figure 12: here, the ‘state exploration’ agent clearly outperforms the ‘parameter exploration’ and ‘random exploration’ agent. Importantly, this illustrates when the context changes randomly, there is no knowledge that could be carried over from one trial to the next. Thus, *active learning*, which focuses on making observations that allow to transfer insights from one trial to the next, will be ineffective. In contrast, *active inference*, which focuses on making observations that allow for precise inference about the current hidden state (context) at a trial, provides an effective solution to this problem (cf., Figure 9), such that this agent always correctly infers the current context of a trial and, in consequence, whether to sample the safe or risky option.

**Figure 13.**
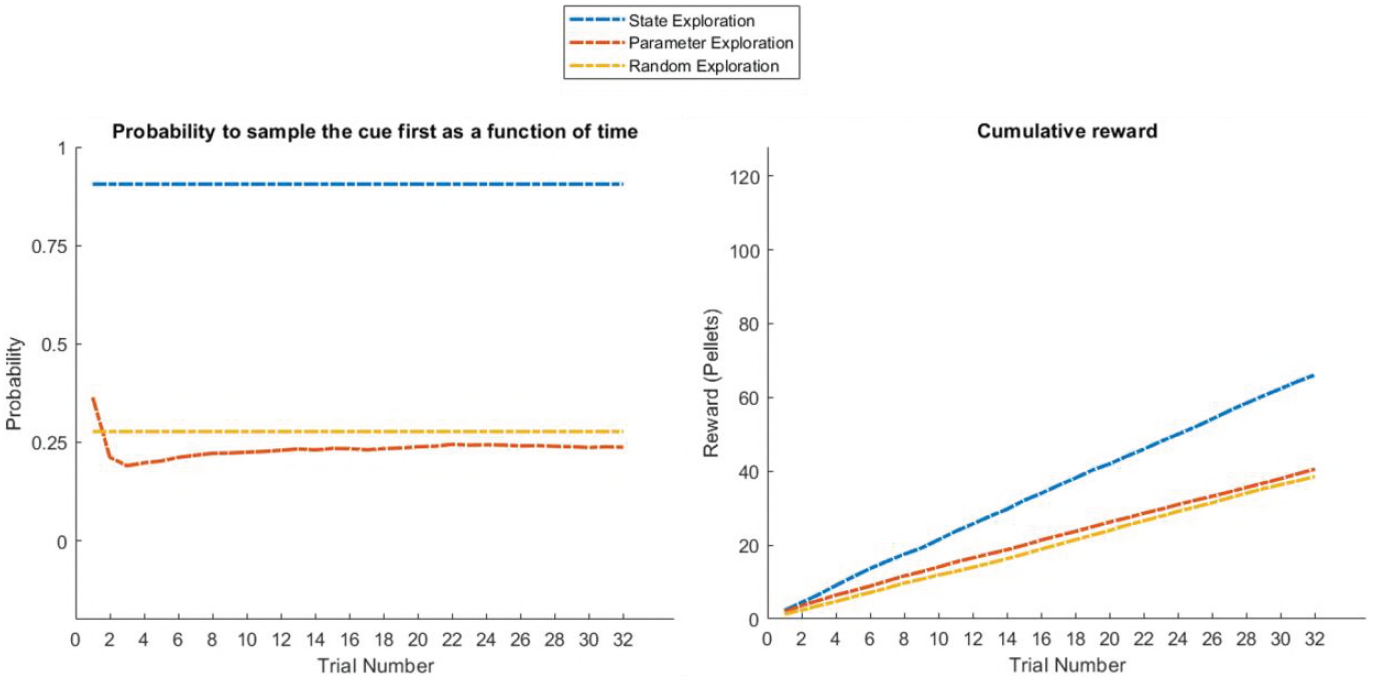
Response profiles of a ‘state exploration’, ‘parameter exploration’ and ‘random exploration’ agent in a task that requires inference. In the problem introduced in Figure 9, where an agent can infer the randomly changing context from a cue, ‘parameter exploration’ will be ineffective, because there is no insight that could be transferred from one trial to the next. ‘State exploration’, in contrast, provides an effective solution to this task, because it allows an agent to infer the current context on a trial-by-trial basis. Left: probability to choose the informative cue at the beginning of a trial. This shows that only the ‘state exploration’ agent correctly infers that it has to sample the cue at the beginning of every trial to adjust its behaviour to the current context (defined as a high or low reward probability in the risky option). Consequently, it outperforms the ‘parameter exploration’ and ‘random exploration’ agent in its cumulative earnings in this task (right).

Figure 14 compares the three agents in the task introduced in Figure 10, which has the same design as the previous example but now with a stable (high or low) reward context across the entire experiment. This task can be solved with both active learning and active inference. The active learning agent has a high bias for sampling the risky option in the beginning of the experiment, and will thus learn whether it is associated with a high or low reward probability. The active inference agent has a strong preference for sampling the cue in the beginning of the experiment, but can adjust its prior over the current context due to stable (high or low reward) feedback from the cue (as illustrated in Figure 10). Thus, both the ‘state exploration’ and ‘parameter exploration’ agent will clearly outperform the ‘random exploration’ agent.

**Figure 14.**
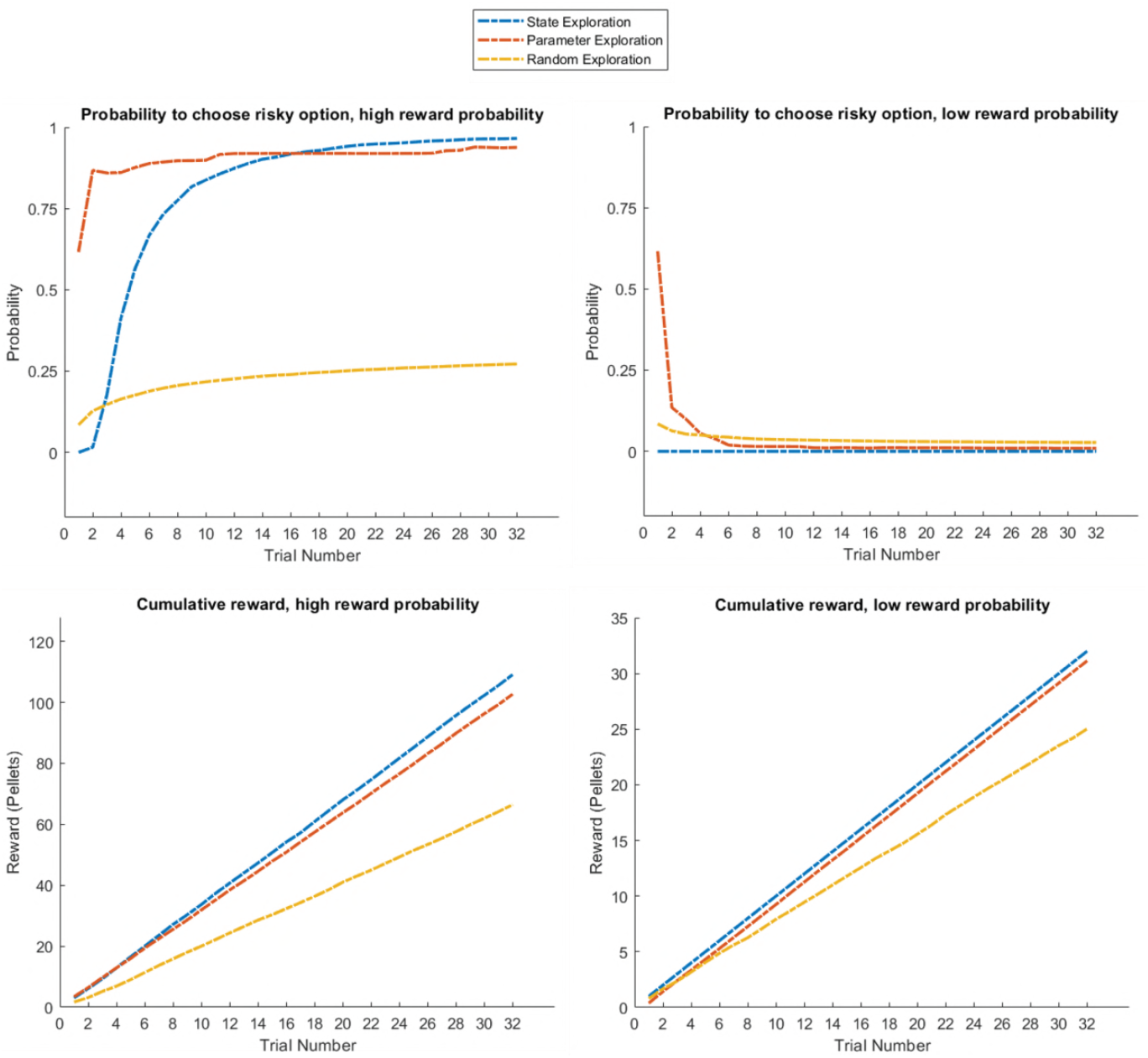
Response profiles of a ‘state exploration’, ‘parameter exploration’ and ‘random exploration’ agent in a task that requires learning or inference. Same problem as in Figure 13, but now with a stable high or low reward context (as in Figure 10). This task can be solved by either sampling the risky option to learn about its reward statistics (‘parameter exploration’), or sampling the cue to learn about the current context and adjusting the prior over contexts due to constant feedback from the cue (‘state estimation’). This can be seen in the response profiles in the upper panel, such that the ‘parameter exploration’ agent has a strong preference for sampling the uncertain risky option in the beginning of the trial (left and right), while the ‘state exploration’ agent only starts sampling the risky option at the beginning of the trial if it has sampled the cue several times before, which always indicates a high reward context (left, cf. Figure 10). This leads to a similar performance level of these two agents as measured by the cumulative reward, which exceeds the performance of the ‘random exploration’ agent (lower panel).

## Conclusion

We have illustrated the emergence of active inference and active learning when casting choice behaviour as probabilistic inference. Under the assumption that behaviour maximises model evidence or (equivalently) minimises surprise over future outcomes, this implies that choice behaviour will reflect a tendency to fulfil preferences and maximise utility, but also to minimise uncertainty about the current state of the environment as well as about relevant task contingencies. Whilst the tendency to fulfil ones preferences reflects exploitative behaviour, uncertainty reduction induces exploratory behaviour. We have contrasted such ‘goal-directed’ exploratory behaviour with ‘random’ exploration caused by imprecise and stochastic behaviour that is unrelated to an agent’s uncertainty about the world.

This perspective makes specific predictions for behaviour. In particular, it introduces a distinction between the uncertainty about current states, which can be resolved by active inference, and uncertainty about model parameters, which can be resolved via active learning. Both uncertainties motivate goal-directed exploratory behaviour but make different predictions for actual decision-making. Minimising the uncertainty over hidden states predicts that agent’s will seek observations from which there is a clear and precise mapping to the underlying hidden state, such as moving to a vantage point to infer the location of prey or sampling a cue that allows to infer the current context, as illustrated in the simulations above. Importantly, this sort of uncertainty reduction depends on a particular representation of the structure of the task and a particular parameterisation of that representation, which allows an agent to assess the mapping from observations to hidden states. We argue that agents are also driven by minimising the uncertainty about this parameterisation itself, as illustrated in the first simulations on ‘parameter exploration’. Minimising the uncertainty over model parameters can even result in behaviour that conflicts with minimising the uncertainty over hidden states – in situations where agents try to sample options that are associated with high ambiguity but also with high novelty or information gain. Consequently, a key prediction for behaviour is that the uncertainty about contingencies will modulate the effect of uncertainty about hidden states on behaviour. An option will be very interesting (i.e. informative) if its outcomes are ambiguous due to high uncertainty about the mapping from this option to possible outcomes, but the same option will be highly aversive if the agent is very certain that it leads to ambiguous outcomes.

In other instances, ‘model parameter exploration’ and ‘hidden state exploration’ can motivate similar types of behaviour. Our simulations, however, highlight an important conceptual distinction between active learning (first example) and active inference (second example) in their respective time courses. As mentioned above, it is possible to cast our ‘hidden state exploration’ example as an active learning problem, if we assume that the current context is stable enough to be learned over time. A key requirement for *learning* the context, however, is that it is possible to carry information from one trial to the next. If this continuity is broken, for example by changing the context randomly on every trial, the agent has to use active inference in order to gain information about the task. Thus, our framework predicts that active learning will be particularly useful if there are stable regularities or rules in the world that can be learned. Active inference, on the other hand, will be useful if behaviour has to adapt to trial-by-trial changes in the world. For example, imagine your favourite craft beer brewery introduces a novel beer based on the flavour of coffee and oranges. This might present a suitable instance for actively learning about the parameterisation of your preferences for coffee and orange flavoured beer, resulting in a large novelty- or curiosity-bonus for this choice. However, you might be aware that you have a strong preference for Lager over Stout. Consequently, before placing your order, it might be useful to actively infer the hidden state of the novel beverage by asking the bartender whether you will receive a Lager or a Stout.

These considerations also highlight the distinction between ‘goal-directed’ exploratory behaviour in the form of minimising uncertainty about hidden states or model parameters, ‘random’ exploratory (i.e., imprecise) behaviour and exploitative decision-making. The trade-off between these behavioural tendencies is governed by their relative precision. For example, if an agent strongly prefers one particular outcome over all other outcomes, she will display predominantly exploitative behaviour with the aim of attaining this outcome. In contrast, if there is one option that is associated with very high uncertainty about its mappings to outcomes, behaviour will be dominated by sampling that option until its associated uncertainty is resolved (as illustrated in our simulations). Our simulations also illustrate that random exploration becomes adaptive if active learning or active inference is broken (or impossible). If the uncertainty about model parameters and hidden states fails to inform behaviour, the only way to learn about the world is through a higher degree of random sampling of different options. Our simulations have shown that this is the only way to (slowly and inefficiently) learn about the advantage of novel options in the absence of goal-directed exploratory behaviour. Further, it is important to note that these types of exploration themselves depend on a model of the task, such as an observation model or a model of the transitions between states. It will be a key challenge for future work to understand how agents build and compare these models in the first place, which provide the basis for inference and learning.

In summary, we have highlighted the distinction between learning about the world as a consequence of random or imprecise behaviour (‘random exploration’) and goal-directed uncertainty reduction. Further, we have shown how these types of behaviour arise when casting behaviour as probabilistic inference. Importantly, we have identified two types of goal-directed exploratory behaviour, namely *active learning* that reduces the uncertainty that relates to the parameterisation of an agent’s generative model of the world, and *active inference* that reduces uncertainty about hidden states in the world given an agent’s generative model. This former type of uncertainty-reduction will compel an agent to sample novel contingencies that enable learning about the true mappings and thus induce ‘model parameter exploration’. The latter type of uncertainty-reduction about hidden states motivates agents to sample salient observations that allow for precise, unambiguous inference about the current state, thus performing ‘hidden state exploration’. We have shown that this distinction makes relevant predictions for the predominance of different types of exploration in different tasks depending on whether active learning or active inference is more adaptive. This will be critical for understanding the different motives underlying curiosity and information-seeking in animals and artificial intelligence, and provides mechanistic insight into suboptimal choice behaviour arising from broken *active inference* or *active learning*.

## Acknowledgements

KJF is funded by the Wellcome Trust (Ref: 088130/Z/09/Z). TUH is supported by a Wellcome Sir Henry Dale Fellowship (211155/Z/18/Z) and grants from the Jacobs Foundation and the Brain & Behavior Research Foundation. The Max Planck UCL Centre is a joint initiative supported by UCL and the Max Planck Society. The Wellcome Centre for Human Neuroimaging is supported by core funding from the Wellcome Trust (203147/Z/16/Z).

## Code Availability

These routines are available as Matlab code in the SPM academic software: http://www.fil.ion.ucl.ac.uk/spm/. The simulations in this paper can be reproduced and customised via a graphical user interface: by typing >> DEM and selecting the **epistemic learning** demo.

